# Transcriptional regulatory logic of the diurnal cycle in the mouse liver

**DOI:** 10.1101/077818

**Authors:** Jonathan Aryeh Sobel, Irina Krier, Teemu Andersin, Sunil Raghav, Donatella Canella, Federica Gilardi, Alexandra Styliani Kalantzi, Guillaume Rey, Benjamin Weger, Frederic Gachon, Matteo Dal Peraro, Nouria Hernandez, Ueli Schibler, Bart Deplancke, Felix Naef

**Affiliations:** The Institute of Bioengineering, School of Life Sciences, Ecole Polytechnique Fédérale de Lausanne, CH-1015, Switzerland; Department of Molecular Biology, University of Geneva, Geneva, CH-1211, Switzerland; Center for Integrative Genomics, Faculty of Biology and Medicine, University of Lausanne, Lausanne, CH-1015, Switzerland; Department of Diabetes and Circadian Rhythms, Nestlé Institute of Health Sciences, CH-1015 Lausanne, Switzerland; School of Life Sciences, Ecole Polytechnique Fédérale de Lausanne, CH-1015, Switzerland; Center for Integrative Genomics, Faculty of Biology and Medicine, University of Lausanne, 1015 Lausanne, Switzerland; Swiss Institute of Bioinformatics, 1015 Lausanne, Switzerland; Bioinformatics Core Facility, Swiss Institute of Bioinformatics, 1015 Lausanne, Switzerland; Department of Oncology and Ludwig Center for Cancer Research, Faculty of Biology and Medicine, University of Lausanne, 1011 Lausanne, Switzerland; Interfaculty Institute of Bioengineering, School of Life Sciences, Ecole polytechnique Fédérale de Lausanne, 1015 Lausanne, Switzerland; Vital IT, Swiss Institute of Bioinformatics, 1015 Lausanne, Switzerland; Bioinformatics and Biostatistics Core Facility, School of Life Sciences, Ecole polytechnique Fédérale de Lausanne, 1015 Lausanne, Switzerland; Department of Molecular Biology, Faculty of Sciences, University of Geneva, 1211 Geneva, Switzerland; Present address: Dualsystems Biotech AG, 8952 Schlieren, Switzerland; Present address: Nestlé Institute of Health Sciences, 1015 Lausanne, Switzerland; Present address: University of Montpellier, 34095 Montpellier, France

## Abstract

Many organisms exhibit temporal rhythms in gene expression that propel diurnal cycles in physiology. In the liver of mammals, these rhythms are controlled by transcription-translation feedback loops of the core circadian clock and by feeding-fasting cycles. To better understand the regulatory interplay between the circadian clock and feeding rhythms, we mapped DNase I hypersensitive sites (DHSs) in mouse liver during a diurnal cycle. The intensity of DNase I cleavages cycled at a substantial fraction of all DHSs, suggesting that DHSs harbor regulatory elements that control rhythmic transcription. Using ChIP-seq, we found that hypersensitivity cycled in phase with RNA polymerase II (Pol II) loading and H3K27ac histone marks. We then combined the DHSs with temporal Pol II profiles in wild-type (WT) and *Bmal1*^*-/-*^ livers to computationally identify transcription factors through which the core clock and feeding-fasting cycles control diurnal rhythms in transcription. While a similar number of mRNAs accumulated rhythmically in *Bmal1*^*-/-*^ compared to WT livers, the amplitudes in *Bmal1*^*-/-*^ were generally lower. The residual rhythms in *Bmal1*^*-/-*^ reflected transcriptional regulators mediating feeding-fasting responses as well as responses to rhythmic systemic signals. Finally, the analysis of DNase I cuts at nucleotide resolution showed dynamically changing footprint consistent with dynamic binding of CLOCK:BMAL1 complexes. Structural modeling suggested that these footprints are driven by a transient hetero-tetramer binding configuration at peak activity. Together, our temporal DNase I mappings allowed us to decipher the global regulation of diurnal transcription rhythms in mouse liver.

## Introduction

Circadian clocks provide mammals with cell-autonomous and organ-based metronomes that relay diurnal environmental cues to temporal gene expression programs [1,2]. In particular, diurnal rhythms in mRNA transcription result from the combined actions of the autonomous circadian oscillator, systemic signals and other temporal cues such as feeding-fasting cycles [3–6]. While it is commonly assumed that around 10% of genes exhibit cyclic mRNA levels in the liver [7], this number increases to nearly 50% when only considering liver-specifically expressed genes [8]. Moreover, these mRNA rhythms cover a continuum of peak times [9,10]. Although mRNAs can also rhythmically accumulate due to post-transcriptional regulation [6,11–14], it is of interest to obtain a more comprehensive view on transcriptional regulators and mechanisms underlying time-specific diurnal transcription. In a light-dark (LD) cycle, two main waves of transcription are found, one during the day (at around ZT10) and the other towards the end of the night (around ZT20), accompanied by dynamic chromatin state modifications [6,11,12].

Current models of time-specific transcription in the liver involve the core clock transcription factors (TFs) BMAL1/CLOCK that activate transcription maximally at ZT6 [15–17], as well as the nuclear receptors RORs and REV-ERBs, whose targets are maximally transcribed around ZT20 [18,19]. Rhythmically active TFs also include clock-controlled outputs, notably the PAR-bZIP proteins (DBP, TEF, HLF), maximally active near ZT12 [16,20]. Furthermore, diurnally fluctuating systemic signals may drive rhythmic TF activities, for example, HSF1 shuttles to the nucleus and activates transcription at ZT14 [21,22], and similarly, SRF shows activity at the night-day transition [23]. Moreover, regulators controlled by feeding-fasting cycles include FOXO TFs that are active during the day, CREB/ATF family members at the light-dark transition, and SREBP during the night [5,24]. Finally, the glucocorticoid receptor (GR) signals the onset of behavioral activity (light-dark transition) [25].

Frequently, these factors act by binding to sequence-specific DNA elements located in the vicinity of gene promoters [26,27], however, less is known about more distally located enhancer regulatory elements involved in diurnal transcriptional control. To start identifying such regulatory elements, recent maps of the activity related chromatin mark H3K27ac[6], as well as enhancer RNAs (eRNAs) [28], were established. These studies identified thousands of putative enhancers with a broad range of peak activity times, which were associated with distinct DNA regulatory motifs and TF binding patterns.

Here we used genome-wide DNase I hypersensitivity mapping [29] to further identify temporally active transcriptional regulatory elements. In the context of the circadian clock, DNase I hypersensitive site mapping was first applied to study regulation of the *Dbp* gene in mouse liver, which led to the identification of several DNase I hypersensitive sites (DHSs) located in 5’-flanking and intronic regions [30]. Several of those regions showed diurnal rhythms in hypersensitivity with amplitudes as large as three fold, which prompted us to generate a temporally resolved and genome-wide DNase I hypersensitivity map [31,32]. We detected around 65’000 DHSs in mouse liver, of which 8% cycled. Moreover, such cycling hypersensitivity occurred in phase with Pol II loadings and H3K27ac histone marks, suggesting that DHSs harbor regulatory elements controlling rhythmic transcription. Analysis of WT and circadian clock deficient *Bmal1*^*-/-*^ animals enabled us to identify transcription regulators with activities at specific times of the day, and to explore how feeding rhythms affect oscillatory activation of transcription in the absence of a functional circadian clock. By contrasting DHS sites in WT and *Bmal1*^*-/-*^ animals, we uncovered that BMAL1 has limited but specific impact on DNA accessibility in regulatory regions. Finally, because DNase I hypersensitivity mapping leaves characteristic footprints at sites of bound TFs, we could study the temporal dynamics of TF complexes bound to DNA. This allowed us to propose a temporal DNA binding mode for the BMAL1:CLOCK hetero-tetramer complex.

## Results

### DNase I hypersensitive site mapping during diurnal cycles in mouse liver

To identify DNA regulatory elements controlling diurnal transcriptional rhythms in the mouse liver, we mapped DHSs every 4 hours during a full light dark (LD) cycle. Specifically, C57BL/6 male mice were kept in standard 12h light, 12h dark cycles, and four animals were sacrificed every 4 hours for one day followed by liver dissection (Methods). DNase I hypersensitivity libraries were produced, sequenced, and mapped to the mouse genome using standard methods (Methods). To monitor transcription activity in the same conditions, we generated ChIP-seq samples for the histone modification H3K27ac (marking active regulatory elements [33]) and re-sequenced previous total Pol II ChiP-seq libraries [12] at increased coverage (Methods and Table S1). Circadian clock outputs result in the rhythmic transcriptional activation of hundreds of genes, notably through binding of CLOCK:BMAL1 heterodimers [16,17]. To validate our assays, we therefore examined the known circadian output gene *Dbp* (Movie S1), maximally transcribed at ZT8 [30], to determine whether cutting frequency at DHSs exhibited diurnal variation. We detected several DHSs in the vicinity of *Dbp*, with high intensity and narrow signals surrounded by low noise levels (Fig 1A). Overall, enriched sites coincided well with regions identified using classical DHS mapping [30], and overlapped with BMAL1 ChIP-seq regions [17] (Fig S1). As exemplified by a DHS nearby the transcription start site (TSS) of *Dbp*, we observed that DHSs were located in regions with lower H3K27ac signals in between H3K27ac-enriched islands, suggestive of TF-induced nucleosome displacement [34–36] (Fig 1B). The DNase I hypersensitivity changed diurnally, notably at the TSS (Fig 1C) where the oscillations in DNase I hypersensitivity, Pol II, and H3K27ac peaked in sync at ZT10 (Fig 1B). Moreover, all DHSs within 15 kb of the *Dbp* TSS displayed oscillations with the same phase as the TSS (Fig 1D), suggesting regulatory relationships between these regions and gene transcription.

**Fig 1.**
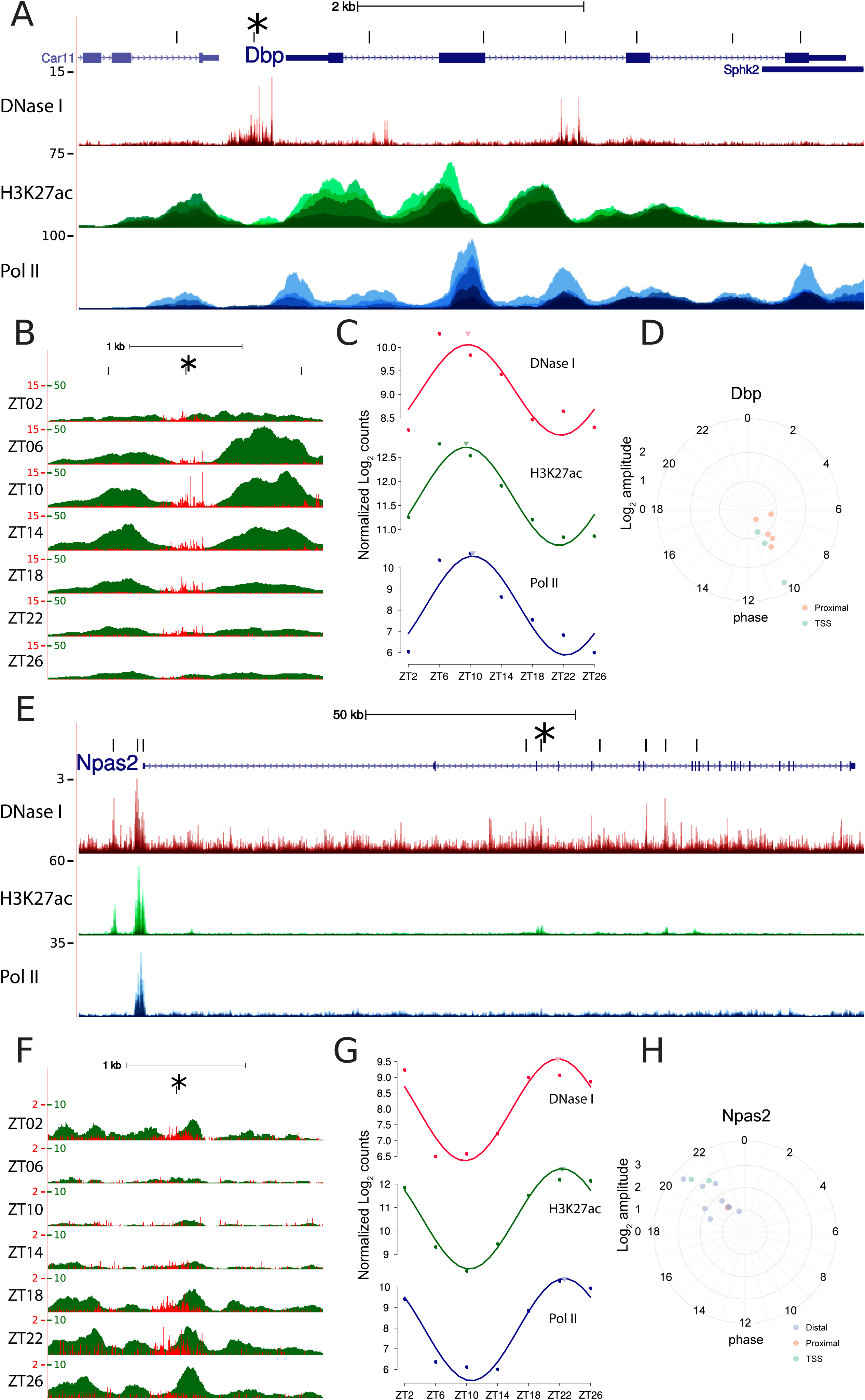
DNase I hypersensitivity is rhythmic during diurnal cycles in mouse liver. A. DNase I hypersensitivity, Pol II density, and H3K27ac enrichment at the *Dbp* locus. The DHS track shows the frequency nucleotide-resolved DNase I cuts, while H3K27ac and Pol II ChIP-seq signals are smoothed over 100 bp. All time points are overlaid. The center of each DHS-enriched region is indicated on top and corresponds exactly with previously identified DHSs (Fig S1). B. Zoom-in around the DHS at the TSS of *Dbp* (position marked with a star also in A) reveals dynamics of DNase I cuts around the clock. Both DNase I and H3K27ac signals are maximal at ZT10 and minimal at ZT22, consistent with BMAL1-mediated activation of *Dbp* transcription. C. Read counts (in log_2_ units) for DNase 1 cuts (in windows of +/- 300 bp) centered on the *Dbp* TSS. Idem for Pol II and H3K27Ac ChIP-seq reads (in windows of +/- 1000 bp) centered on the same DHS and cosine fits show a common peak time around ZT10. Peak to trough amplitudes are about 16-fold for Pol II, and approximately 4-fold for both DNase I and H3K27ac. D. Phases and amplitudes of all DHS sites located in the neighborhood of the *Dbp* gene (nearest-TSS association according to annotation). Distances from the center of the plot indicate log_2_-amplitudes, and angles (clockwise from ZT0) indicate peak times. We observed that all regions oscillate around a common phase of ZT10. E.-H. Similar to A-D but for *Npas2*, which has an opposite phase to *Dbp*, i.e. it peaks near ZT22.

We next analyzed the *Npas2* gene (Movie S2), another known clock target [37]. *Npas2* is a target of RORs and peaks in the late night-time around ZT22 [38]. We detected several DHSs along the transcribed region of *Npas2* (Fig 1E), including proximal (defined as 1-10kb from a TSS) and distal (defined as >10kb from a TSS) sites. The distal sites displayed high amplitude oscillations of DNase I signals and H3K27ac (Fig 1F). Normalized signals at the *Npas2* TSS also peaked at the expected phase with maximal signal at ZT22 for all three marks studied (Fig 1G). Finally, all DHSs associated with *Npas2* (those having *Npas2* as their closest TSS), including numerous distal regions, likewise cycled with phases around ZT22 (Fig 1H). The examples of the the *Dbp* and *Npas2* loci suggest that our genome-wide study detected DHSs with high resolution, and that the temporal patterns of DNase I cuts reflected diurnal activities of these elements.

### Identification of regulatory elements and transcription factor footprints in mouse liver DHSs

To comprehensively map putative regulatory elements genome-wide, we merged our DNase I hypersensitivity time points and performed peak finding (Methods). This revealed 62’418 DHS sites, covering around 2% of the mappable genome (considering a width of 600 bp for each DHS site), which is comparable to previous studies across mouse tissues [39] (all sites and associated signals in Table S2). Because we aimed at associating DHSs with nearby genes to infer regulatory relationships, we first discarded transcripts from ENSEMBL annotations that were not expressed in our samples. For this, we used histone modifications, Pol II profiles, and now also DNase I signals at transcription start and end sites of annotated transcripts to train a supervised learning method (support vector machine) that distinguishes expressed (active) from non-expressed genes, similar to our previous work [12] (Methods). To infer putative regulatory relationships, we then annotated each DHS to the nearest active TSS. Distances between DHSs and TSSs followed a bimodal distribution, with a first mode around 100 bp from the TSSs and a second 10 kb from the TSS (Fig S2A). Consistent with previous reports [40,41], one third of our DHSs were found within 1kb of TSS, while almost half were located more than 10kb from a TSS (Fig S2B). At TSSs, the genomic distributions of DNase I cuts, Pol II, and H3K27ac signals (centered on TSSs) were consistent with accessibility of DNA being determined by nucleosome displacement and Pol II complex assembly (Fig S2C) [42]. At distal DHSs, profiles of H3K27ac showed a dip in the peak center, consistent with occupation by TFs and nucleosome displacement (Fig S2D), while the weaker Pol II signals could reflect distal assembly of the transcriptional complex [43], or interactions between enhancer regions and the TSS through DNA looping [44,45].

To determine whether DHSs reflected DNA-bound transcription regulators, we searched for short windows protected from cleavage, or footprints [46] within a +/- 300 bp window around the center of each DHS. This identified previously reported footprints, as illustrated for the well-characterized promoter of the *Albumin (Alb)* gene [47] (Fig S2E). In the promoter region of *Reverb*α *(Nr1d1)*, the detected footprints coincided with E-boxes and high BMAL1 ChIP-seq signals (Fig 2A). Overall, the majority (70%) of DHSs within 1 kb of a TSS contained at least one footprint, while this proportion dropped to one half for proximal (defined as DHSs within 1-10kb of a TSS) or distal (>10kb of a TSS) DHSs (Fig 2B). Since transcribed DNA is known to be DNase I sensitive [48], the DHSs without footprints might reflect transcription. To test this, we analyzed the number of footprints in DHSs outside of promoter regions and further marked with H3K36me3, a mark coinciding with transcribed gene bodies [12,49]. Indeed, DNase I hypersensitive regions without footprints were frequently (90%) linked with highly transcribed genes (Fig 2C). Thus, DHSs at TSS seemed to contain more footprints than distal DHSs, and transcription elongation explains why some DNase I hypersensitive regions did not exhibit a footprint.

**Fig 2.**
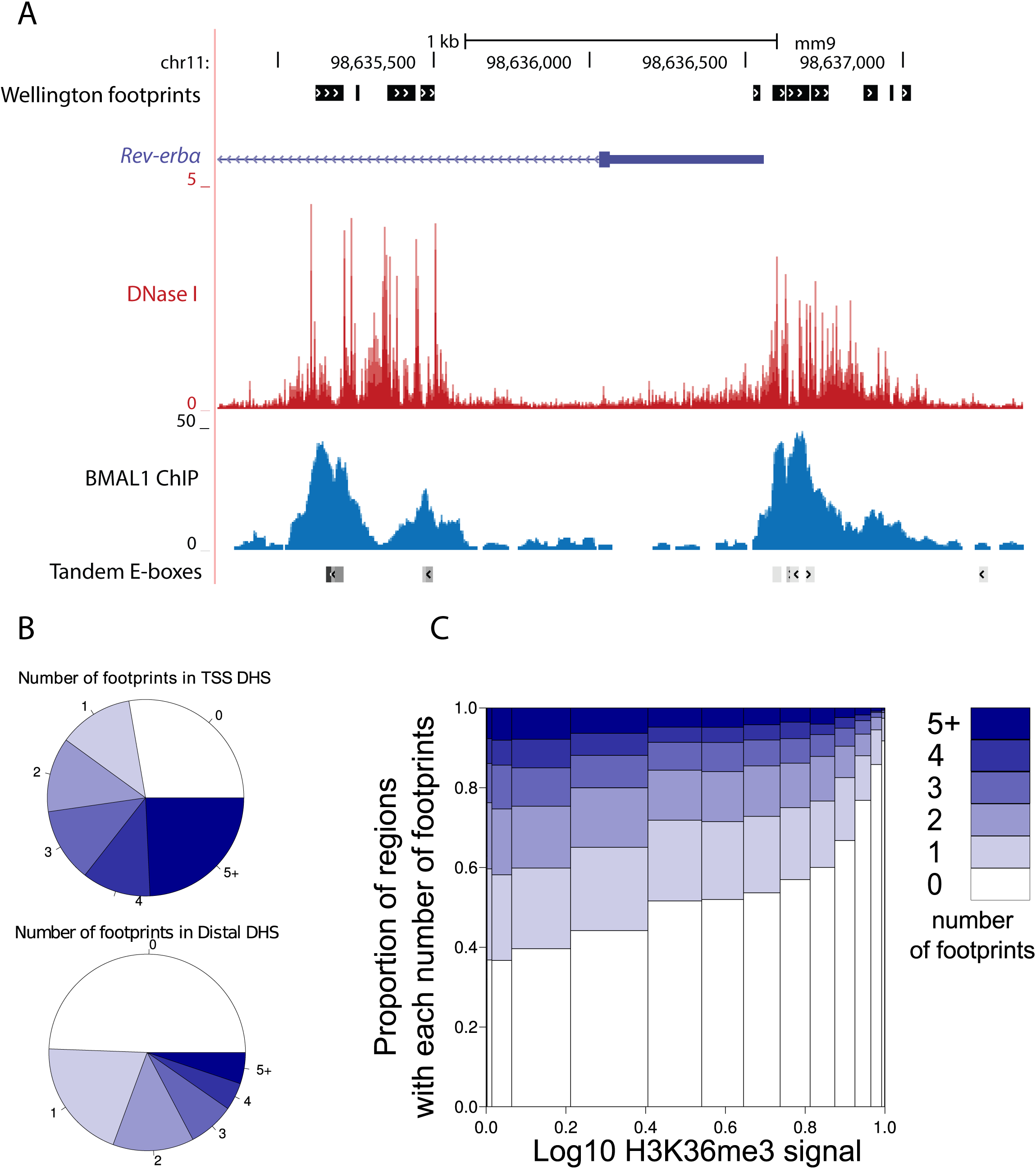
Location-dependent footprint characteristics of DHSs. A. Visualization of DNase I signal (red) around the *Rev-erb*α promoter with the footprints (detected by Wellington) annotated in black, on top. This region contains BMAL1 binding sites (blue) with E-box motifs, annotated on the bottom line, which is marked by a characteristic footprint. The DNase I cleavage pattern is lower at the binding site, reflecting protection of the DNA from digestion, whereas high signals are observed on the edges of the binding site. B. Number of footprints within DHSs (+/- 300 bp around the peak center). TSS regions contain more footprints on average. More than half of distal regions contain a footprint. C. Number of footprints detected in DHSs in function of (relative) H3K36me3 signal.

### TSSs and distal regulatory elements display 24-hour oscillations in DNase I hypersensitivity in sync with Pol II and H3K27ac enrichment

We next studied whether DNase I hypersensitivity, Pol II density, and H3K27ac quantified at the identified DHSs displayed diurnal rhythms using harmonic regression (Methods). The number of cyclic regions identified at different significance thresholds clearly indicated that Pol II and H3K27ac oscillated at a larger number of DHSs compared to the DNase I signal itself, both for proximal and more distal DHSs (Fig 3A). To select rhythmically active regions, we assessed the combined rhythms of the three marks at each DHS as previously using Fisher’s combined test [12,50], which yielded 4606 DHSs (7.3%, FDR<0.05). For all three signals, the amplitude of the oscillations was larger at distal DHSs (the median peak-to-trough amplitude was two-fold for DNase I and H3K27ac, and higher for Pol II) compared to TSSs, and Pol II had larger amplitudes than either DNase I or H3K27ac (Fig 3B). Moreover, the peak times of the oscillations in DNase I signals were, except for some small deviations, similarly distributed as peak times in gene transcription and H3K27ac [6,11,12], with a weak evening peak around ZT10 and a marked late night peak around ZT22 (Fig 3C). We next considered the relationships of peak times in the DNase I, Pol II and H3K27ac rhythms. It is known that many chromatin marks exhibit diurnal rhythms that are tied to transcription [6,11,12,16], and similarly, enhancer RNAs (eRNAs) were shown to be transcribed in sync with their cognate transcripts [28]. We observed that DNase I cuts, Pol II, and H3K27ac displayed synchronous oscillations at DHSs (Fig 3D). Such relationships were maintained after removing DNase I sensitive regions situated in the transcribed region of active genes (Fig S3), indicating that this phenomenon was not a mere reflection of transcription [51]. To test whether the signals measured at DHSs near TSSs were temporally correlated with those at proximal or distal DHSs, we examined pairs of oscillating DHSs (FDR <0.1, Fisher’s combined test), of which one was located near a TSS (<1kb) and the other in an intergenic region positioned at least 2kb and at most 20kb from any TSS. While no pair reached statistical significance for DNase I signals (at the level of FDR<0.1), probably reflecting that DNase I signals are noisier than the two other marks, we found 1611 pairs oscillating for H3K27ac and 630 for Pol II. The two peak times were highly correlated with a differences within one hour (Fig 3E), suggestive of enhancer-TSS interactions [40].

**Fig 3.**
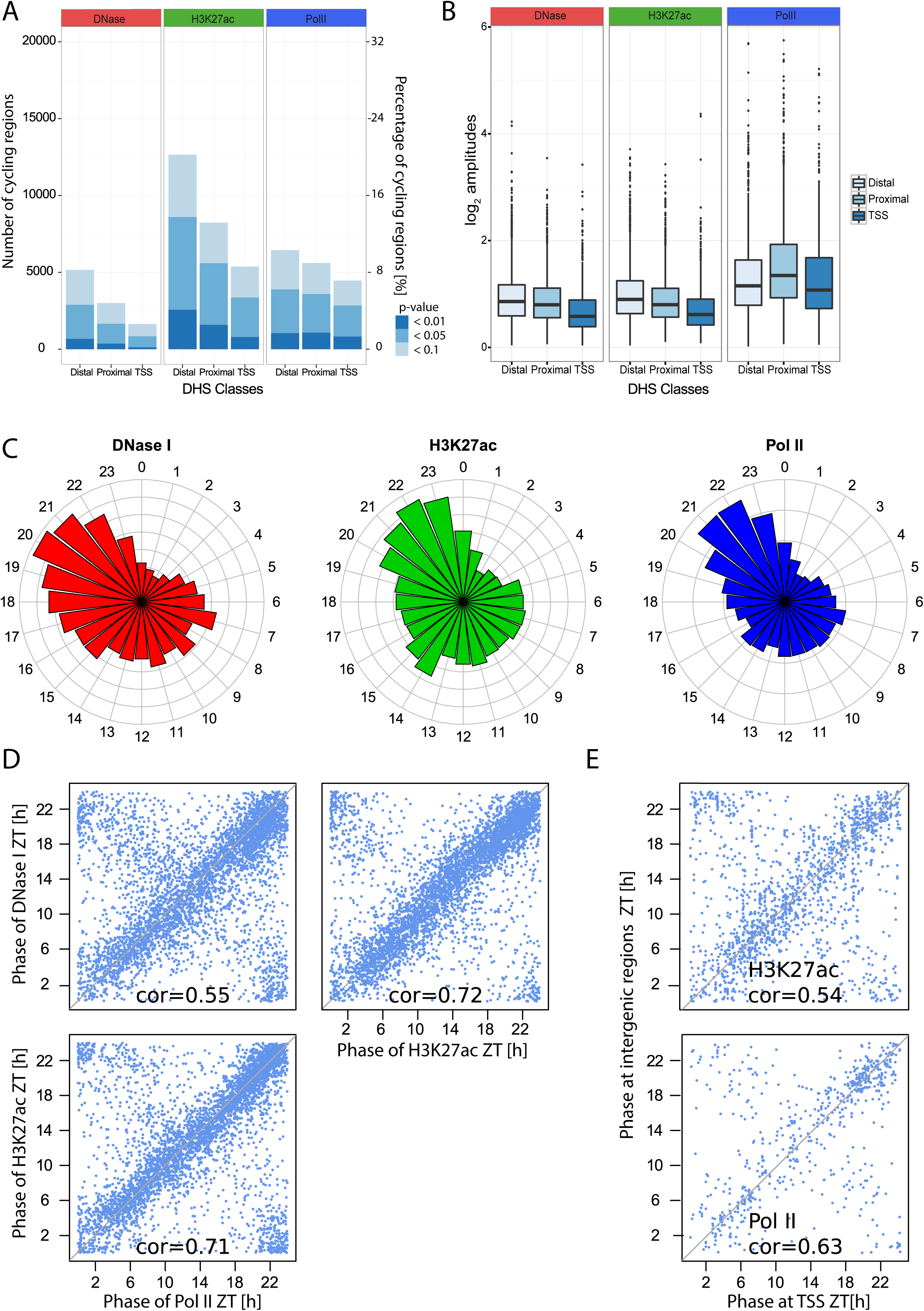
Genome-wide rhythms in DNase I signals are synchronous with Pol II transcription and histone acetylation. A. Number of DHSs with statistically significant cycling DNase I signals (left), H3K27ac signals (middle), or Pol II signals (right) at three different thresholds (p < 0.1, p <0.05 and p <0.01, harmonic regression), partitioned according to their genomic location: TSS (1 kb), proximal (1-10 kb from TSS), or distal (>10 kb from TSS). B. Comparison of log_2_ amplitudes for DHSs in each class (TSS, proximal and distal) and in each signal (Pol II, H3K27ac and DNase I). 4609 sites were selected (FDR<0.05, Fisher’s combined test). Higher amplitudes were observed in distal and proximal regions compared to TSSs (p < 2.2*10^−16^, t-test). In addition, Pol II loadings showed higher peak-to-trough ratios than the two other signals. C. Circular histograms representing the distributions of phases for each mark at DHSs selected as in B. D. Comparisons of peak times between DNase I, Pol II and H3K27ac at DHSs (DHSs selected with p < 0.05, Fisher’s combined test), diagonals are indicated in gray. Values of circular correlations are indicated (p < 10^−10^, circular correlation). E. Relationships of peak times between DHSs in intergenic regions with their nearest TSS (pairs selected with FDR<0.1, Fisher’s combined test). 1611, respectively 630 significant pairs were found for H3K27ac and Pol II signals.

### Computational analysis identifies transcription factors through which the circadian clock and feeding-fasting cycles control diurnal gene expression

To understand how the circadian clock and the feeding-fasting cycle control diurnal gene expression in liver, we studied mRNA expression and Pol II loading at TSSs in WT and *Bmal1*^*-/-*^ mice subject to the same, night restricted, feeding regimen (Fig S4). First, we observed that a similar number of genes oscillated in the WT and *Bmal1*^*-/-*^ genotypes (p < 0.05), however, with an overlap of about 30% for Pol II and 50% for mRNA. This indicates that genes with a diurnal expression differ between WT and *Bmal1*^*-/-*^ mice (Fig S4A). While such comparisons are based on cutoffs, stratifying by peak-to-trough amplitudes clearly showed that high amplitude rhythms are more abundant in WT as compared to *Bmal1*^*-/-*^ mice (Fig S4B), and that this was more pronounced for mRNA than for Pol II loading at TSSs. For example, we found twelve genes with greater than ten-fold mRNA amplitudes in WT, and only three in *Bmal1*^*-/-*^ mice. Genes with Pol II or mRNA rhythms in both genotypes showed highly correlated phases, with a tendency for a slight average delay (~1 hour in Pol II and less in mRNA) in the absence of a circadian clock (Fig S4C).

Functional annotation using KEGG and Reactome pathways and comparison between mRNA rhythms in WT and *Bmal1*^*-/-*^ animals showed that genes annotated for circadian rhythm as well as lipid and sugar metabolism were enriched in the WT condition. In *Bmal1*^*-/-*^ mice, we observed that pathways related to sugar and lipid metabolism were still oscillating, notably SREBP and ChREBP signaling (Table S3). To identify transcriptional regulators underlying rhythmic transcription by the circadian clock and feeding-fasting cycles, we used a computational approach that combines temporal Pol II loading at TSSs in WT and *Bmal1*^*-/-*^ mice with annotated transcription factor binding sites in accessible chromatin regions as defined by our DHSs. Using DHSs and a collection of about 1900 position-weight matrices for TF-DNA affinities (Methods), we identified DNA sequence motifs that explain rhythmic Pol II patterns in WT and *Bmal1*^*-/-*^ mice. Briefly, we modified previously described linear models [17,52,53] to identify transcriptional activities (strictly speaking DNA motifs) represented by phase (time of maximal activity) and amplitude (Methods). In this model, motif activities are linearly combined, as in the phase vector model [54], according to the presence of corresponding DNA motifs within nearby DHSs. This enabled us to take into account, in addition to the proximal promoter, a collection of putative regulatory regions that may control the expression of a given gene (Fig 4A). Specifically, we considered motifs in DHSs located within a certain window from active promoters, and first estimated the optimal window size according to the quality of the fit. We found that the inclusion of DHSs up to 50kbp was improving the fits in both genotypes (Fig 4B), suggesting that enhancers (represented by distal DHSs) contribute to circadian gene transcription. In WT mice (Fig 4C, Table S4), our modeling confirmed that known circadian transcription factors showed the strongest activities, as reflected by the emergence of ROR responsive elements (RREs) [18,55] with predicted peak activity at ZT22, D-Box elements at ZT12 [56], and E-boxes around ZT8, as previously described [57]. Other motifs that had previously been associated with diurnal transcription in the liver were also identified. These included Forkhead box (FOX) motifs around ZT20 and ZT5 [58–62], the CREB motifs at ZT7 [63–67], GR motifs around ZT10 [68], SREBP motifs at ZT19 [24,69,70], HSF1 at ZT16 [21,22,26], and ETS TFs in the morning [28]. In *Bmal1*^*-/-*^ mice (Fig 4D, Table S4), activities of E-Box, RRE, and D-Box motifs were not detected or greatly reduced, as expected in the absence of a functional circadian oscillator. On the other hand, transcription factors linked with metabolic functions, notably those associated with feeding rhythms (e.g. FOX, CREB, SREBP) were identified among the strongest contributors in the absence of a functional clock. Similarly, transcription factors whose activity depends on systemic signals (e.g. GR and HSF1) were also found with peak activity times that were similar in the WT and *Bmal1*^*-/-*^ mice. Interestingly, CREB was found among the most delayed transcription factor activities, with a predicted delay of six hours (Table S4). To test this prediction, we measured nuclear levels of CREB and pCREB using Western blots of nuclear extract from four independent livers every two hours in WT and *Bmal1*^*-/-*^ mice (Fig 4E and Fig S5). On average, we observed a phase delay of approximately two hours in *Bmal1*^*-/-*^ mice. Although this was not significant (p=0.5, Chow test), presumably owing to inter-individual variability in the feeding patterns, it is consistent with predictions by our model. Of note, similar inter-individual variability has been reported for the rhythmic activation of the TORC1 and AMPK pathways [71].

**Fig 4.**
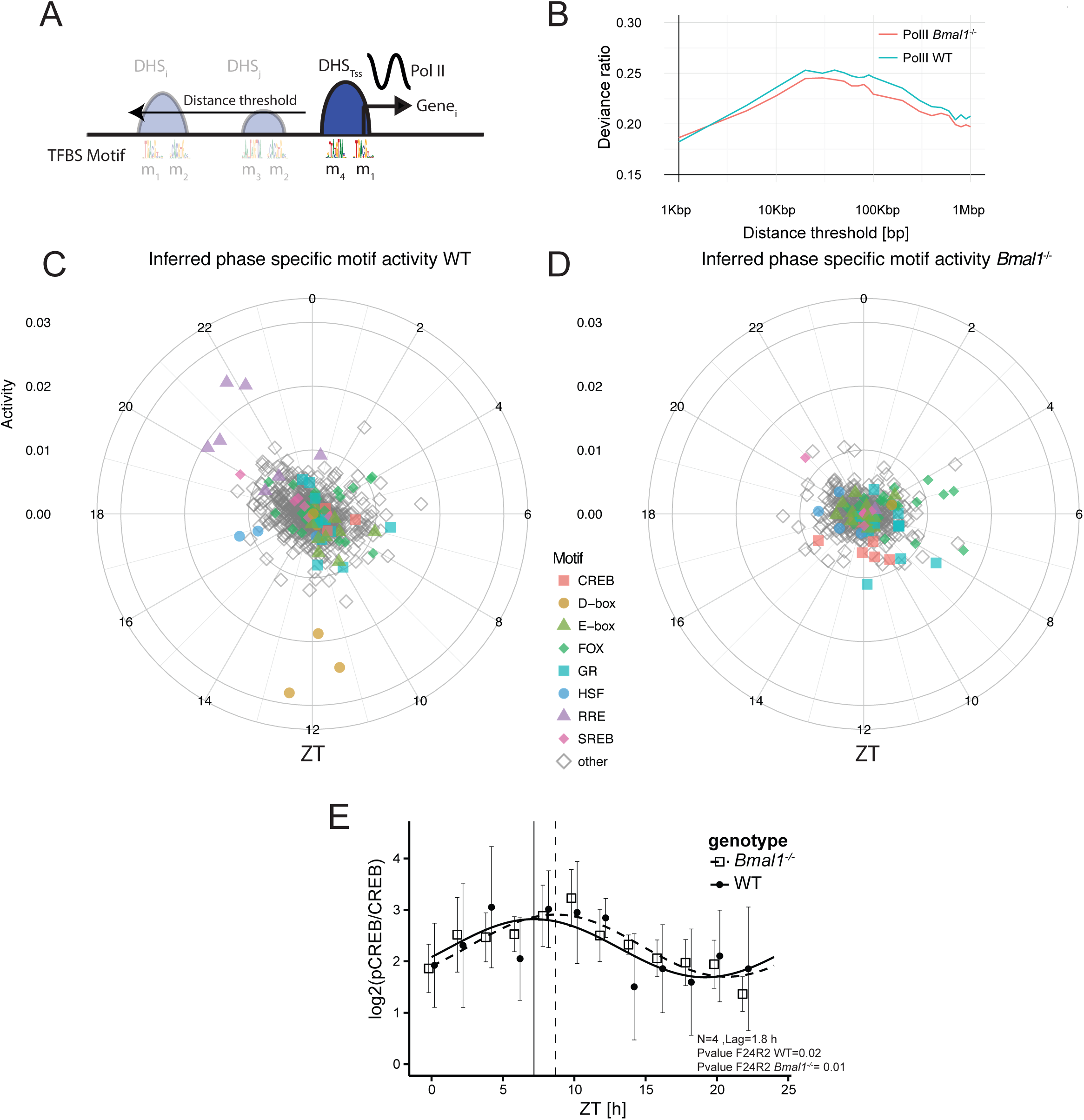
Distal DHSs help identify diurnally active transcription regulators. A. Scheme of the linear model to infer active transcription regulators: TF motifs in DHSs within a symmetric window around active TSSs are used to explain diurnal rhythms in transcription. B. Fraction of explained temporal variance (Deviance ratio) in Pol II loading (at the TSS of all actives genes) for WT and *Bmal1*^*-/-*^, in function of the window size (radius) for DHS inclusion, shows a maximum at around 50 kb. Here *α*=0 was used in the glmnet (Methods). C.-D. Inferred TF motif activities for WT and in *Bmal1*^*-/-*^ shown with amplitudes (distance from center) and peak times (clockwise, ZT0 at the top), using a window size of of 50 kb. All 819 (WT) and 629 (*Bmal1*^*-/-*^) motifs (overlap is 427) with non-zero activities are shown. Note though that most activities are very small and cluster in the center. Certain families of TFs are indicated in colors (full results are provided in Table S4). Radial scale for activities is arbitrary but comparable in C and D. E. Quantification of Western blots for pCREB and CREB in WT and *Bmal1*^*-/-*^ genotypes (log_2_ (pCREB/CREB)). Nuclear extracts from four independent livers were harvested every two hours. Both genotypes showed a significant oscillation (p<0.05, harmonic regression) of the mean signal from the four mice. Though the peak time in *Bmal1*^*-/-*^ mice is delayed by 1.8 hours, the comparison of the rhythm in the two genotypes is not significant (p=0.49, Chow test). Individual blots are presented on Fig S4.

### BMAL1 has specific impact on DNA accessibility in regulatory regions

We next examined how BMAL1 binding might influence DNA accessibility. In our *Bmal1*^*-/-*^ mice [72], we performed DHS mapping at ZT6, near the maximal DNA binding activity of BMAL1 in WT mice. DNase I hypersensitivity at BMAL1-bound sites (detected in ChIP-seq) [17], such as in the *Rev-erb*α locus, was markedly decreased in *Bmal1*^*-/-*^ mice, whereas control (unbound) regions like the *Gsk3* promoter showed no difference (Fig 5A). Overall, we observed a clear shift in DNase I hypersensitivity at DHSs with BMAL1 binding sites. Regions bound by BMAL1 in the WT [17] showed fewer DNase I cuts in *Bmal1*^*-/-*^ as compared to WT animals, indicating that BMAL1 binding specifically impacts DNA accessibility at its target sites (Fig 5B). These findings are consistent with the proposed pioneering function of the BMAL1-CLOCK complex [36]. While DNase I signals at those sites were also significantly lower at minimal BMAL1 activity in the WT (ZT18), the *Bmal1*^*-/-*^ mice showed even lower signals (Fig 5C). The same analysis at sites bound by the E-box binding protein USF1 [73] did not show such differences between WT and *Bmal1*^*-/-*^ animals (Figs 5D and 5E).

**Fig 5:**
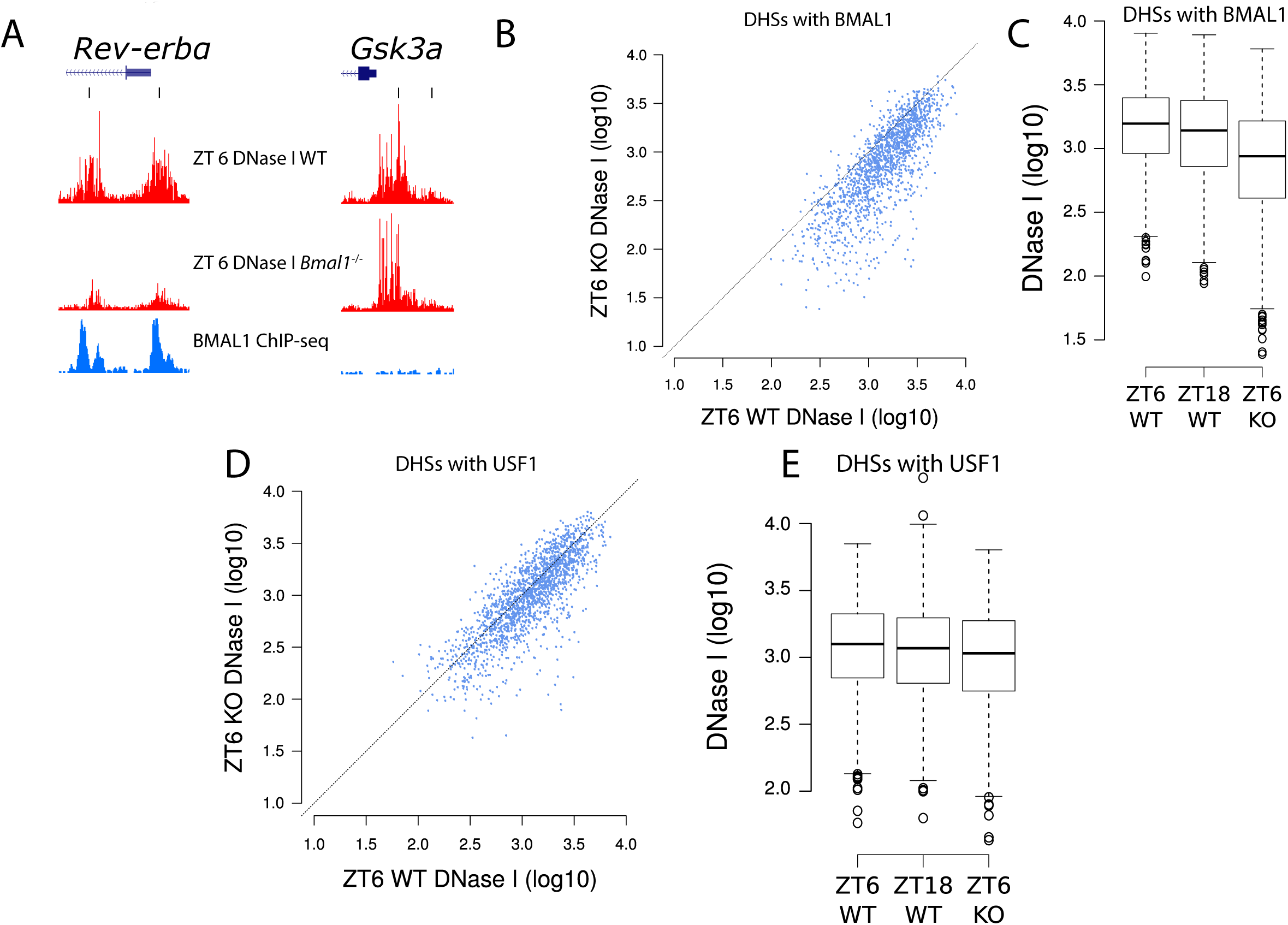
Chromatin accessibility is generally similar in *Bmal1*^*-/-*^ and wild-type mice, but lower at BMAL1 bound sites in the former. A. The *Rev-erb*α (left) and *Gsk3a* (right) promoters, where DHSs are indicated with black ticks at the top. DNase I signal (in red) is strongly reduced in *Bmal1*^*-/-*^ mice at sites bound by BMAL1:CLOCK in WT mice (BMAL1 ChIP-seq signal in blue) in the *Rev-erb*α promoter, but similar in WT and *Bmal1*^*-/-*^ mice at the *Gsk3a* promoter, not bound by BMAL1. The vertical scale is the same for all four DNase I tracks, as well as for both BMAL1 ChiP-seq tracks. B. Comparison of DNase I signals at ZT6 in *Bmal1*^*-/-*^ versus WT mice. All DHSs overlapping BMAL1 ChIP-seq peaks in [17] are shown (n=1555). C. Boxplots showing DNase I intensity at the same sites as in B, at peak (ZT6) and trough (ZT18) activities of BMAL1 in the WT, and at ZT6 in *Bmal1*^*-/-*^ mice. D.-E Same as B-C, but using overlap with USF1 ChIP-seq peaks [73] to select DHSs (n=1705).

### DNase I footprints at BMAL1 sites reveal temporal exchanges of transcription factor complexes

Owing to the 3D structures of protein-DNA interactions, genomic patterns of DNase I cleavage around transcription factor binding sites display factor-specific footprints [32,74–77]. We previously showed that BMAL1 binds DNA rhythmically, and that strong BMAL1 binding was frequently associated with tandem E-boxes [78] separated by 6 or 7 nucleotides, which were bound by one or two BMAL1/CLOCK dimers [17]. Here, we analyzed DNAse I footprints at BMAL1 binding sites as a function of time. Starting from BMAL1 ChIP-seq sites, we modified a “mixture model” for DNase I cuts [79] to determine the optimal boundaries of the footprints at each time point, as well as the probability that the factor is bound to DNA (calculated here as the probability that the DNase I showed a footprint) for every site (Methods). We then analyzed footprints at BMAL1 binding sites containing tandem E-boxes separated by 6 bp (E1E2-sp6). At ZT6, close to the maximal DNA binding activity of BMAL1, both E-boxes in the E1E2-sp6 motif appeared to be protected from digestion. In contrast, at ZT18 only the 5’ E-box displayed a footprint consistent with occupation by a transcription factor (Fig 6A, full time course in Fig S6). Moreover, the footprint at ZT18 was undistinguishable from that in the *Bmal1*^*-/-*^ mice, suggesting that other transcription factors bind BMAL1 sites when BMAL1 activity is low. The estimated proportion of E1E2-sp6 motifs showing a footprint indicative of two BMAL1/CLOCK dimers varied across time points, with a maximum of 65% at ZT10, and minimum of 20% in the *Bmal1*^*-/-*^ animals (Fig 6B). Also, the binding dynamics of BMAL1 at E1-E2-sp7 (tandem E-boxes separated by 7bp) was largely similar to that for E1-E2-sp6, though E1-E2-sp7 had both E-boxes predominantly protected only at ZT6, suggesting spacer-specific binding dynamics (Fig S7). In contrast, the footprints at BMAL1 binding sites with single E-boxes did not show significant changes in time or in the *Bmal1*^*-/-*^ mice (Fig S8), again suggesting that other bHLH transcription factors can also bind at BMAL1 sites. In fact, footprints at DNA regions bound by the bHLH transcription factor USF1 in ChIP-seq [73] were largely similar to that of BMAL1 sites with single E-boxes, though the fraction of sites with clear footprints was reduced for USF1 compared to BMAL1 (Fig S9).

**Fig 6:**
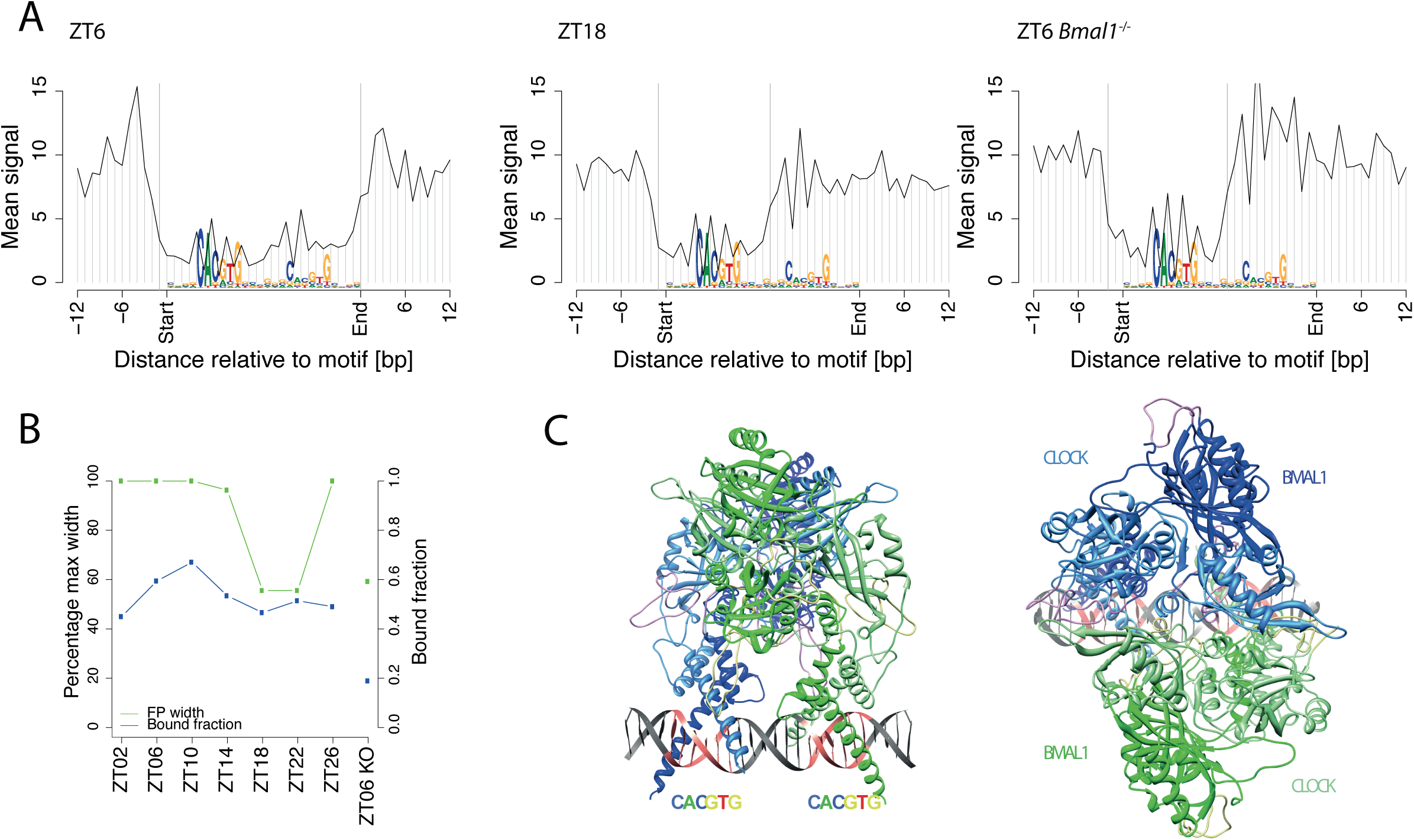
BMAL1 footprints indicate temporally changing protein-DNA complexes, consistent with binding of a hetero-tetramer to DNA. A. Genomic profiles of DNase I cuts around double E-boxes with a spacer of 6 bp (E1-E2 sp6). We selected n=249 E1-E2 sp6 motifs overlapping a BMAL1 ChIP-seq peak, and show the average of profiles for loci classified as bound by the mixture model (posterior probability > 0.5). At ZT6, we observed that nucleotides around both E-boxes are protected. In contrast, at ZT18, the width of the protected region is reduced by approximately half, with the second E-box no longer protected from digestion. The signals are anchored to the motif position. Orientation of sites and signals is according to the best match to the E1-E2 sp6 motif. In *Bmal1*^*-/-*^, only one E-box appears occupied. B. Width (left-side y-axis, green) of the protected region in WT and in *Bmal1*^*-/-*^, for E1-E2 sp6 motifs occupied by BMAL1. Fraction of predicted occupied sites is shown in blue (right-side y-axis). C. Two views of the 3D computational model of the CLOCK:BMAL1 hetero-tetramer showing two heterodimers of CLOCK:BMAL1 occupying an E1-E2 sp6 site. The two heterodimers are shown in green and blue, while darker green and darker blue correspond to BMAL1 and lighter colors to CLOCK proteins. Information content along the DNA strands is shown in grey with highly constrained nucleotides of the motif in red.

To better understand the time-dependent footprint at BMAL1 sites and to gain insight into how the CLOCK:BMAL1 heterodimer occupies its tandem E-box-containing target sites, we used recently established 3D protein structures of single BMAL1/CLOCK complexes combined with molecular modeling (Methods). Our models strongly support formation of CLOCK:BMAL1 heterodimers in a hetero-tetramer configuration at peak activity of these factors, and residual binding of the dimer or other transcription factors during low activity times. Two 3D models of the hetero-tetramer configuration were constructed. In the first model, the spacing between the two E-boxes was 6 bp (sp6) (Fig 6C, Movie S3, Fig S6, File S2) and in the second model the spacing was 7 bp (sp7) (Movie S4, Fig S7). For the model of the single CLOCK:BMAL1 complex, we used the crystal structure of the heterodimeric CLOCK:BMAL1 (pdb id: 4F3L) [80], into which we built the missing parts of the flexible loops. To link the single CLOCK:BMAL1 model to the E-box, we employed the complex crystal structure of CLOCK:BMAL1 basic helix-loop-helix domains bonded on the E-box (CACGTG) (pdb id: 4H10) [81]. We then superimposed the two single CLOCK:BMAL1 E-box models, with the sp6 DNA and the sp7 DNA, forming the respective symmetric hetero-tetramer models. We found that the 6 bp spacing between the two E-Boxes was optimal to establish favorable interactions between the two CLOCK:BMAL1 heterodimers, involving mainly residues (e.g., K335, Y338, Q352, E380 and E384) located in the PAS-B domain of the CLOCK in a dynamic H-bond network [82]. Similarly, the 7 bp spacing seemed also able to favor a hetero-tetramer conformation, producing only a minor twist of 10° in the three interval base pairs. However, a conformation with base pair spacing less than 6 or more than 7 would make complex formation difficult because of conformational constraints. Thus, the modeling results are consistent with two CLOCK:BMAL1 heterodimers binding binding to two E boxes separated by 6 or 7 base pairs, and the DNase I footprints with characteristic and dynamically changing shapes suggest exchanges of different transcription factor complexes on the DNA during the diurnal cycle.

Finally, we examined temporal footprints at DHSs bound by other rhythmically active TFs, REV-ERB, HSF1, SREBP and CREB (Fig S10). Interestingly and unlike what we observed for BMAL1/CLOCK, the shapes of the footprints for those factors did not change with time and were unaffected in the absence of BMAL1. However, the fraction of sites showing footprints coincided well with the maximal transcriptional activity of the different factors. For example, footprints centered on REV-ERBα-bound ROR response elements (RREs) showed the largest proportion of footprints at ZT22, which coincides with the trough activity of the REV-ERB repressors. The low percentage of bound (as detected in ChIP) RREs with footprints called by the model was low (<20%), which could reflect that nuclear receptors tend to have a low residence time and therefore display a lower DNase I cleavage-protection pattern [76]. For HSF1, the number of footprints was maximal at ZT18, approximately four hours later than the previously reported peak activity [83], and for the feeding-induced SREBP this number peaked during the night, as expected [24]. Lastly, high confidence CREB binding sites [84] showed clearly marked and invariable width footprints throughout the 24 hours in both WT and *Bmal1*^*-/-*^ mice, consistent with the finding that CREB activity is regulated post-translationally on the DNA [63–66].

## Discussion

### DNase I hypersensitivity shows daily rhythms in adult mouse liver in sync with transcription and chromatin activity marks

We mapped genome-wide DNase I hypersensitivity with 4-hour time resolution in the livers of adult mice. This provided a comprehensive view on the dynamics of chromatin accessibility controlled by the circadian clock, feeding/fasting cycles, or both. Overall, the identified hypersensitive regions, clustered in about 60’000 DHSs (typically several hundreds of base pairs wide and covered two percent of the mappable genome. One third of these regions was located near gene promoters and the remaining two thirds more distal from TSSs, which is consistent with what has been previously observed in mammalian cells [40]. On a genome-wide scale, 98’000 footprints were detected in about 60% of these accessible regions. Importantly, our data provided global insights into the temporal variations in DNase I hypersensitivity on the timescale of several hours to one day. Indeed, while it was previously shown that high amplitude circadian genes such as *Dbp* showed nearby hypersensitive regions [30], it was not known how widespread these rhythms are genome-wide. Here, we showed that thousands of DHSs exhibit rhythmic signals with peak-to-trough amplitudes that are comparable to those of Pol II signals. Accessible chromatin, as measured with DNase I hypersensitivity, is typically associated with transcriptionally active states, and often reflects the presence of proteins bound to regulatory DNA elements [31,32,40]. However, we showed that DNase I sensitive regions within gene bodies, notably in the case of highly transcribed genes, may reflect transcription elongation. As a consequence, they do not necessarily display DNA footprints such as the ones discerned in regulatory elements.

We then compared the temporal patterns of DNase I hypersensitivity with other frequently used transcriptional activity marks, in particular H3K27ac and Pol II. While DNase I signals, H3K27ac and Pol II densities all showed abundant rhythmicity, H3K27ac abundance cycled at the largest number of DHSs, in particular at distally located sites. For both DNase I hypersensitivity and H3K27ac, the peak-to-trough amplitudes appeared higher in distal elements as compared to TSSs. Such dynamic accessibility might reflect increased protein binding dynamics at enhancers, suggesting their potential role in controlling diurnal gene expression. This would parallel mechanisms underlying cell type specificity, where the modulation of histone marks and accessibility of chromatin at enhancers are among the major features associated with regulatory mechanisms [85]. The hypothesis that distal DHSs might represent enhancers for diurnal transcription was further supported by our observation that rhythms in pairs of putative enhancers and nearby TSSs showed a tight temporal correlation. In contrast to the observed delay between H3K4me3 enrichment and Pol II density, reported previously [12], no significant delays were observed between accessibility as measured by DNase I hypersensitivity and H3K27ac enrichment. This likely reflects that turnover of histone acetylation is faster than that of histone methylation [86]. We then used these temporal datasets to explore the involvement of putative enhancer regions in the cyclic recruitment of Pol II at the TSSs and subsequent transcription of the respective target genes. Our findings were consistent with a previous study on enhancer RNA (eRNA), which showed that eRNAs cluster in specific circadian phases and are correlated with Pol II occupancy and histone acetylation [6,28]. In addition, eRNA levels are correlated with the expression of nearby genes [28].

### BMAL1 knockout animals subjected to a nighttime-feeding regimen show abundant Pol II and mRNA rhythms

Our genome-wide study of Pol II loading and mRNA expression in WT and *Bmal1*^*-/-*^ mice kept under LD cycles and night-restricted feeding revealed that the number of genes exhibiting diurnal fluctuations did not drastically change in these behaviorally arrhythmic animals. However, we found that the number of high amplitude oscillations in mRNA accumulation was much reduced in *Bmal1*^*-/-*^ mice. All in all, our observations suggest that feeding cycles can entrain a significant set of low amplitude transcriptional oscillations, while the circadian clock drives high amplitude rhythms of a relatively limited number of transcripts (Fig S4).

### Combining DHSs with genomic sequence can predict transcription factors with cycling activities in the presence or absence of BMAL1

In this study, we accumulated compelling evidence for the contribution of distal regulatory elements in circadian transcription regulation. In fact, we observed that about 47% of DHS are located at more than 10 Kb from the closest active TSS. Using penalized regression models, we predicted a collection of transcription factor binding motifs that best explain diurnal variation in transcriptional activity in both WT and *Bmal1*^*-/-*^ mice. Moreover, while the analysis of promoter sequences recently yielded insights into promoter architecture that favor rhythmic transcription [87], the inclusion of distal DHSs up to 50 kb improved the variance explained by our penalized linear model in WT and *Bmal1*^*-/-*^. The obtained set of transcription factors that exhibited high activity amplitudes in WT was similar to the one derived from a screen that used differential display of DNA-binding proteins [21]. On the other hand, comparison with *Bmal1*^*-/-*^ mice indicated that transcription regulators related to feeding/fasting cycles and rhythmic systemic signals were active in both genotypes, as would be expected. Among those, Forkhead domain factors (FOX) have been implicated in cell cycle regulation and oxidative stress, and are negatively regulated by insulin signaling [59]. Notably, FOXO1 and FOXO6, like the core clock [4], regulate the expression of key enzymes implicated in gluconeogenesis [60,88], collectively pointing towards FOX transcription factors as effectors of metabolic rhythms in liver.

We also found CREB to be among the most delayed transcription factor activities inferred by the generalized linear model in *Bmal1*^*-/-*^ mouse liver. CREB is implicated in the nutrient response cycle and it regulates hepatic gluconeogenesis [62,63,68,89]. We were able to replicate the pattern of CREB activity, as measured by its phosphorylation on Ser 133, in WT mice [89] and we showed that CREB activity is still oscillating in *Bmal1*^*-/-*^ mice. Thus, our results confirm that CREB is regulated by food-related signaling in clock-deficient mice subjected to a night-restricted feeding regimen. The phase delay of two hours thereby suggests that the circadian clock is implicated in the fine-tuning of hepatic glucose metabolism. Consistently, CREB activity during fasting was shown to be modulated by CRY1 and CRY2, which are rhythmically expressed in the liver [89]. Similarly, transcription factors that are responsive to systemic signals, such as heat shock transcription factor 1 (HSF1) driving rhythmic transcription of heat-shock proteins around ZT18 [21,22], or the glucocorticoid receptor (GR) sensitive to glucocorticoid hormones (GCs) released near the day-night transition [25,90–93], were identified both in WT and *Bmal1*^*-/-*^ mice. Our identification of GR activity is consistent with the previous observation that hundreds of circadian transcripts, distinct from clock-controlled circadian genes, are under glucocorticoid control [94].

### Transcription factor binding dynamically reshapes DNA footprints

Comparing DNase I signals between WT and *Bmal1*^*-/-*^ samples at ZT6 revealed that the majority of BMAL1 binding sites showed a decrease in DHS signals (Fig 5), which may be consistent with a pioneering function for the BMAL1:CLOCK core clock transcription factor [36]. Moreover, our analysis of DNase I signals at nucleotide resolution revealed interesting dynamics in the shape of the footprint, which was reminiscent of our earlier proposition that strong and functional BMAL1:CLOCK recognition elements, as those found near the majority of core circadian clock genes, involved the binding of a dimer of CLOCK:BMAL1 heterodimers [17,73,78,82]. Here, we found that CLOCK:BMAL1 binding leaves a wide footprint spanning a tandem E-box element at the maximum activity, and that this footprint shrinks to encompass only a single E-box at the minimum activity time point, resembling that detected in *Bmal1*^*-/-*^ mice. This indicates that other E-box binding transcription factors expressed in the liver, such as USF1, can occupy these E-box sites. These transcription factors may thereby function as placeholders to render these sites quickly accessible for CLOCK:BMAL1 heterodimers at the onset of the next circadian cycle. Indeed, USF1 has been shown to act as a non-allelic suppressor in certain mouse strains carrying a semi-dominant mutation of CLOCK [73]. Structural modeling of the TF–DNA complexes based on the CLOCK:BMAL1 crystal structures supported the establishment of a hetero-tetramer configuration at peak activity [17,73,78,82].

## Conclusion

We performed temporally resolved DNase I hypersensitivity mapping to identify regulatory elements and transcription factor footprints underlying rhythmic transcription during diurnal cycles in the mouse liver. Our study sheds light on the interrelationships between the nutrient response cycle and the circadian clock as well as the contribution of the distal regulatory elements to circadian gene expression. In sum, we found that hypersensitivity at both promoter proximal and distal sites oscillates in phase with transcription during diurnal cycles. Computational integration of DHSs with transcription activity allowed us to highlight differences in the transcriptional regulatory logic of diurnal cycles in WT and circadian clock-deficient *Bmal1*^*-/-*^ animals. Finally, digital footprint analysis revealed dynamically changing transcription factor complexes on DNA.

## Materials and Methods

### Animals

C57/BL6 male and *Bmal1*^*-/-*^ mice [72] 12–14-wk-old (at time of sacrifice) were housed in a 12 hours light/12 hours dark (LD) regimen. These were then entrained to a 12 hours/12 hours LD regimen with water *ad libitum* but food access only between ZT12 and ZT24 for Pol II ChIP-seq and H3K27ac ChIP-seq and microarray experiments for 7 days (ZT, Zeitgeber time; ZT0 is defined as the time when the lights are turned on and ZT12 as the time when lights are turned off) before the animals were sacrificed. Mice used for DNase I-seq were entrained to a 12 hours/12 hours LD regimen with water and food ad libitum. At each ZT2, ZT06, ZT10, ZT14, ZT18, ZT22, and ZT26, 3-5 mice were anesthetized with isoflurane and decapitated. The livers were perfused with 2 ml of PBS through the spleen and immediately collected. A small piece of liver tissue (approx. 100 mg) was snap-frozen in liquid nitrogen and kept at −80°C for RNA extraction. The remaining liver tissue was immediately homogenized in PBS containing 1% formaldehyde for chromatin preparation. All animal care and handling was performed according to the Canton of Geneva (Ueli Schibler) and Canton of Vaud (Nouria Hernandez and Fred Gachon) laws for animal protection.

### DNase I-seq

Mouse liver nuclei were prepared as described in [95]. Freshly prepared nuclei were suspended in ice-cold Ψ-buffer (11 mM KPO_4_ pH 7.4, 108 mM KCl, 22 mM NaCl, 5mM MgCl, 1 mM CaCl_2_, 1 mM DTT) and pelleted. 5×10^6^ nuclei were suspended in 200 μl of Ψ-buffer supplemented with 0.2% of NP40 and 1 u/ml of DNase I (DPFF Worthington Biochemical Corporation). DNase I digestion was performed for 6 minutes at room temperature and the reaction was stopped by adding 200 μl of lysis buffer (50mM Tris-HCl pH 8, 20 mM EDTA, 1% SDS, 200 μg/ml proteinase K). Protease digestion was performed overnight at 55 °C. RNaseA (100 μg/ml) was then added and samples were incubated at 37°C for an hour. DNA was then extracted twice with phenol-chloroform and precipitated with isopropanol in the presence of 0.5 M NaCl. DNAs were dissolved in 5 mM Tris-HCl pH 8. DNAs from 4 animals were pooled, and 75 μg of DNA were loaded on 11 ml 10%-50% sucrose gradient in STE buffer (1M NaCl, 20 mM Tris-HCl pH 8, 5 mM EDTA) and centrifuged at 30000 rpm for 16 hours at 20°C (SW 40 Ti rotor, Beckman Coulter Inc). The sucrose gradients were then fractionated, and DNA was precipitated by two volumes of ethanol in the presence of 5 μg of glycogen. Fractions containing DNA sized around 300bp were pooled and used for Illumina library preparation.

### ChIP-seq of RNA Polymerase II

For *Bmal1*^*-/-*^ animals, perfused livers were processed for chromatin preparation as described in [16]. The chromatin samples from the five mice were then pooled, frozen in liquid nitrogen, and stored at -80°C. For the ChIP experiments, the following antibodies were used: anti-RPB2 (Santa Cruz Biotechnology, sc-673-18). To determine the optimal amounts of each antibody, we performed pilot ChIP assays and determined the enrichment for a set of promoters by real-time qPCR according to [16]. A total of 1 ml of each chromatin suspension (containing about 60 µg of DNA) was incubated with 10 µg of anti-RPB2, in buffer A (20 mM Tris/HCl (pH 7.5), 150 mM NaCl, 2 mM EDTA) overnight at 4°C on a rotating wheel. 10 µl of protein A bead suspension (25% slurry in buffer A), pre-blocked with 10 µg/ml of salmon sperm DNA and BSA at 4°C overnight, was then added and the incubation was continued for 1 h at room temperature on a rotating wheel. The beads were then washed with dialysis buffer and ChIP wash buffer as described in [96]. Protein–DNA complexes were eluted from the beads, de-cross-linked, and treated with RNase A and, subsequently, with proteinase K, as described in [16]. The DNA concentration was determined by fluorometry on the Qubit system (Invitrogen). A total of 10–12 ng DNA were used for the preparation of the library. Libraries for ultra-high throughput sequencing were prepared with the ChIP-Seq DNA sample kit (Illumina) as recommended by the manufacturer.

### ChIP-seq H3K27ac

For WT and *Bmal1*^*-/-*^ animals, H3K27ac ChIPs were performed according to the method described by [97] with a few modifications. The 100 µl chromatin aliquots was used for each IP and diluted with 900 µl of RIPA buffer (1% NP-40, 0.5% sodium deoxycholate, 0.1% SDS in PBS at pH 7.4) and added to Dynal magnetic beads conjugated with Sheep-antimouse IgG dynabeads (Invitrogen, Cat no: 110-31) pre-treated with 3 µl of polyclonal antibody for H3K27ac (Active motif, Cat no: 39135) for immunoprecipitation of specific complexes. The samples were incubated overnight at 4°C on rotator, then magnetic beads washed 7 times with lithium chloride wash buffer (100 mM Tris at pH 7.5, 500 mM LiCl, 1% NP-40 and 1% sodiumdeoxycholate) and once with 1X TE buffer (10mM Tris-HCl at pH 7.5, 0.1 mM Na_2_EDTA). The chromatin complex was eluted using elution buffer (1% SDS, 0.1 M NaHCO_3_) for 1 h at 65°C using Eppendorf thermo-mixer. The chromatin was then de-crosslinked overnight at 65°C and ChIP DNA purified using Qiagen PCR purification kit and eluted in 50 µl of elution buffer. For qPCR reaction 1.5 µl of 1/10 diluted ChIP DNA was used. Libraries for ultra-high throughput sequencing were prepared with the ChIP-Seq DNA sample kit (Illumina) as recommended by the manufacturer.

### ChIP-seq of HSF1

ChIP-seq of HSF1 was performed according to the method described by [17]. The HSF1 polyclonal antibody was from Stressgen (Enzo Life Sciences, ADI-SPA-901). For each IP 5 μl of HSF1 antibody was used with 250 μl of pre-cleared chromatin. A ChIP library was prepared using 4 independent ChIP experiment at ZT14 and one lane was sequenced to obtain about 20 million uniquely mapped reads.

### CREB and pCREB Western Blot on Nuclear extract

Hepatic nuclear proteins were prepared as described in [27] using the NaCl-Urea-NP40 (NUN) procedure. 10 µg of the nuclear protein extracts were fractionated on an SDS-PAGE and transferred to a PVDF membrane for Western blot analysis. Antibodies against CREB (Chemicon # AB3006) and Phospho-CREB (pSer133) (Chemicon #AB3442) were used at 1:1000 dilutions. Membranes were stained with naphtol blue black in order to quantify the protein loading.

### ChIP-seq and DNase I-seq data analysis

At each time point, DNA sequenced reads were mapped to the mouse genome (*Mus musculus* NCBIM37 genome assembly (mm9; July 2007)) using bowtie through the HTS station portal (available at http://htsstation.epfl.ch) [98]. Duplicate reads were kept to avoid saturation due to high coverage of Hi-seq libraries and DNase I specificities. Quality controls, including the percentage of reads within enriched regions, indicated high overall enrichment at all time points, as about 50% of DNase-seq reads mapped to 1.3% of the genome considered to be accessible (Table S1). Peak calling was done using ChIP-peak [99] (http://ccg.vital-it.ch/chipseq/chip_peak.php) on DNase I signals merged from all ZT time points with the parameters: cutoff=100, vicinity=400, window size=600, threshold=1000. After peak calling, DNase I, Pol II and H3K27ac signals were quantified at each time-point within a window of +/- 300 bp around every peak center (+/-1kb for H3K27ac). The values thus obtained were quantile normalized between time points for each mark.

### Detection of active transcript

Using ChIP-seq data for Pol II, H3K4me3, H3K36me3 and H3K27ac from [12] in the WT condition, a support vector machine classifier (SVM) was used to detect active transcripts among all Ensembl annotated transcript (version NCBIM37). We selected regions of interests to be +/-300 bp around the TSS for Pol II and H3k4me3, idem and also +/-300 bp around the TES for DNase I, and the last 600 bp of each transcript for the gene body mark H3K36me3. Read counts on the same strand as the transcript annotation was counted per 10 bp and quantile-normalized across time. For training, a set of active and inactive transcripts were extracted consisting in the top 10% and bottom 10% respectively, as determined by Pol II RPKM along each transcript. An SVM was trained on these active versus inactive transcripts, and subsequently applied to all transcripts at each time point. Cross-validation indicated that the SVM had satisfactory False Positive and False Negative results for very high or very low Pol II signals (98% of test transcripts were correctly classified either active or inactive at ZT10). Transcripts shorter than 600bp were set to “active” if they had higher Pol II RPKM than the lower quartile of active transcripts. Transcripts were considered active when they were classified as active at minimally one time point. The active transcripts list was used to associate DHS with the closest active transcription start site (TSS). The annotation result provided 13’457 unique active genes linked with at least one DHS (Table S3).

### Rhythmicity analysis and selections of DHSs

Rhythmicity analysis was done as previously[12] using harmonic regression. Throughout (DHS signals, ChIP signals, mRNA expression) log_2_ normalized signals were used in the harmonic regressions. The Fisher combined probability test [50] for Pol II, H3K27ac and DNase I signals was computed to select rhythmic DHSs. This uses a Chi-squared distribution with 2k (k=3 marks) degrees of freedom. The resulting p-value was used to estimate False Discovery Rates (FDR) via the linear step-up method. mRNA microarray in WT and in *Bmal1*^*-/-*^ mice from [12] were reanalyzed using harmonic regression.

### Analysis of published ChIP-seq data

Published datasets of ChIP-seq of CREB [63], USF1 [73], REV-ERBα [100] in the mouse liver were quantified in our DHSs. ChIP-seq experiments such as SREBP [24] and BMAL1 [17] were included. Z-scores were computed for each ChIP-seq in each DHS. Z-scores greater than 2 were used for subsequent footprint analysis in Fig 6.

### Footprint detection in DHSs

Footprints in DHSs were detected using Wellington (pyDNase library) [46] with parameters: -sh 20,36,5 -fdr 0.05 on all DNase samples concatenated. To analyze the shape of footprints, we extended a mixture model for DNase I cuts [79] to determine the optimal boundaries of the footprints at each time point, as well as the probability that the factor is bound to DNA (calculated here as the probability that the DNase I showed a footprint) for every site (Details in File S1).

### Linear model for inference of phase specific motif activities

To identify rhythmic TF activities from temporal Pol II data, we adapted existing methods based on linear regression [101] to the circadian context [54]. Specifically, we estimated transcription factor motif activities *A*_*f*_ by fitting the following linear model:

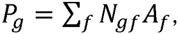

where *p*_*g*_ denotes the 24 hour component of the temporal Pol II profiles, for gene *g*, i.e. *p*_*g*_ = 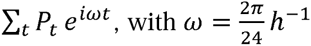. In practice, to perform linear regression with real numbers, we used real and imaginary parts as two dimensional vectors. The matrix *N*_*gf*_ represents the susceptibility of gene *g* to the factor *f*, and contains the motif content for factor *f* in all DHSs within a certain window of an active TSS (Fig 4A). To cover a large representation of TF motifs, we used FIMO [102] and scanned our DHSs using sets of position weight matrices (PWM), from JASPAR [103], TRANSFAC [104], SELEX [105] and WANG [106], in total ~1900 matrices. We counted all motifs below a threshold of 10^−4^. The fitting was performed using the Elastic-net penalized linear regression model [101], which conveniently controls sparseness of the solution (in virtue of the L1 norm), grouping of redundant features (owing to the L2 term, this was important here, since we have a large and redundant set of matrices), and overfitting (using cross-validation). This method is available as an R package called GLMNET and uses the elastic-net penalized regression. Unless otherwise state, we used an ‘alpha’ (tunes the relative weights of the L1 and L2 penalties) value of 0.1. In Fig 4 C-D, real and imaginary components of the inferred activities *A*_*f*_ are plotted, showing both their amplitudes and peak activity times (phases).

### 3D structures of BMAL1/CLOCK heterotetramer

For the single CLOCK:BMAL1, the crystal structure of the heterodimeric CLOCK:BMAL1 (pdb id: 4F3L) was used as an initial model [80]. In this structure there are 5 flexible loops lacking density. The residues in positions 129-134 (length 6 residues), 212-237 (26 residues), 257-275 (19 residues), 291-309 (19 residues) were missing from BMAL1, and the residues 224-247 (24 residues) were missing from CLOCK. These missing parts were computed by Rosetta’s loop modeling application (v3.5); an application that extensively remodels the backbone of the loops [107]. The loops were remodeled and refined by the CCD (Cyclic Coordinate Descent) algorithm [108]. The fragment files, used by CCD were made by Robetta Server [109]. The CLOCK:BMAL1 structure, as a unique chain, was used as Rosetta input and from the output we selected the lowest energy loops for the single CLOCK:BMAL1 model. In order to bind the single CLOCK:BMAL1 model to the E-box, the complex crystal structure of CLOCK:BMAL1 basic helix-loop-helix domains bonded on the E-box (CACGTG) (pdb id: 4H10) was used [81]. This structure was superimposed to the single CLOCK:BMAL1 model with the UCSF Chimera visualization program (v1.5.3) [110]•. In accordance to this super-position the single CLOCK:BMAL1 model the N-terminal helices of CLOCK and BMAL1 was replaced by the helices in the 4H10 structure from the protein data bank. The base-pair geometry of the DNA in the 4H10 structure was analyzed by the 3DNA software (v2.0) [111]••. Two double-strand DNA models, spacing 6 (sp6) and spacing 7 (sp7), with sequence 5’- CACGTGAAAAAA(A)CACGTG-3’, were generated by 3DNA. The CACGTG parts were rebuilt based on the analysis of the DNA in the 4H10 structure. The spacer of 6 bp was built with the standard B-DNA backbone conformation for A-T pairs. For the final models two CLOCK:BMAL1:E-box models were bound to the DNA models with a spacer of 6 bp (sp6) or 7bp (sp7), by superimposing them with UCSF Chimera. In the sp6 model we performed energy minimization for 12500 steps with the NAMD simulation package v2.9. The model was parameterized by the AMBER force field (ff99bsc0) [112].

## Data visualization

Wig files were generated using the bam2wig script [98] and were normalized by the number of mapped reads divided by 10^7^. DNase I signal is represented using the first position of the read alignments considered as the cutting position and without shifting strands. Pol II and H3K27ac are represented using the coverage by the whole read length after shifting forward (in the read orientation) by 80bp and 90bp for, respectively, Pol II and H3K27ac. These wig files were then visualized on the UCSC genome browser (http://genome.ucsc.edu/).

## Data Availability

High-seq Illumina sequencing data for the ChIP-seq (Pol II WT/*Bmal1*^*-/-*^, H3k27ac WT/*Bmal1*^*-/-*^, HSF1 (WT ZT14), and DNase I-seq (WT and ZT6 *Bmal1*^*-/-*^) are available at GEO as the super series GSE60430.

---

**Fig S1.**
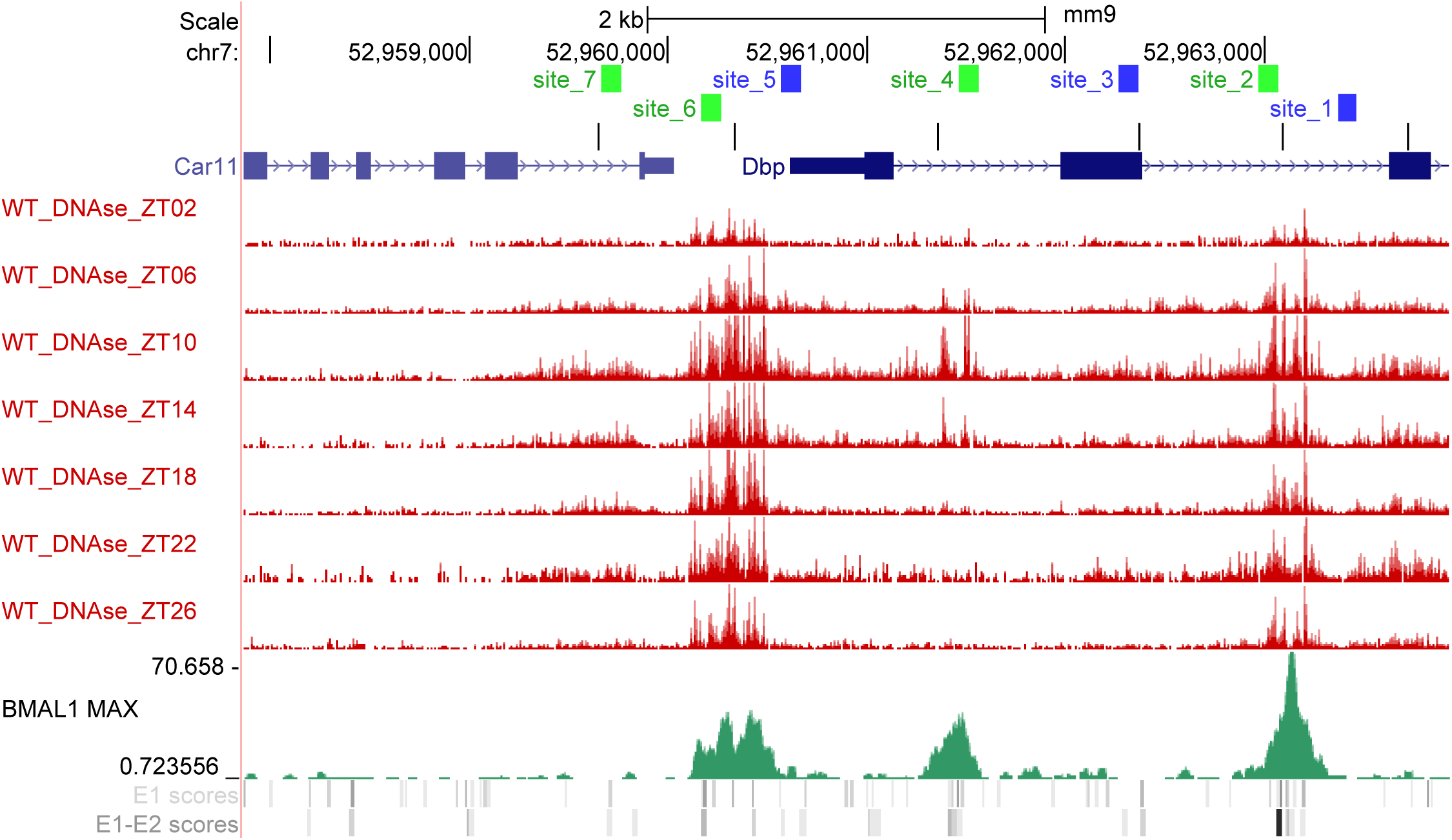
Measured DNase I-seq signals near the *Dbp* gene, compared with previously reporter DHSs in a reference study [30] (marked site_1 to site_7). [30] found seven hypersensitive sites while we detected six DHSs using our peak calling at compatible locations (black marks). Moreover, [30] reported high (sites 2, 4, 6, and 7, in green), or lower (sites 1, 3 and 5 in blue), amplitudes in rhythmic DNase I digestion efficiency, consistent with the DNase I-seq signals (visual inspection). Sites 2, 4, and 7 contain E-boxes that are binding sites for CLOCK and BMAL1. Locations of BMAL1 ChIP-seq signals (bottom track) [17] clearly overlaps strongest DNase I peaks.

**Fig S2.**
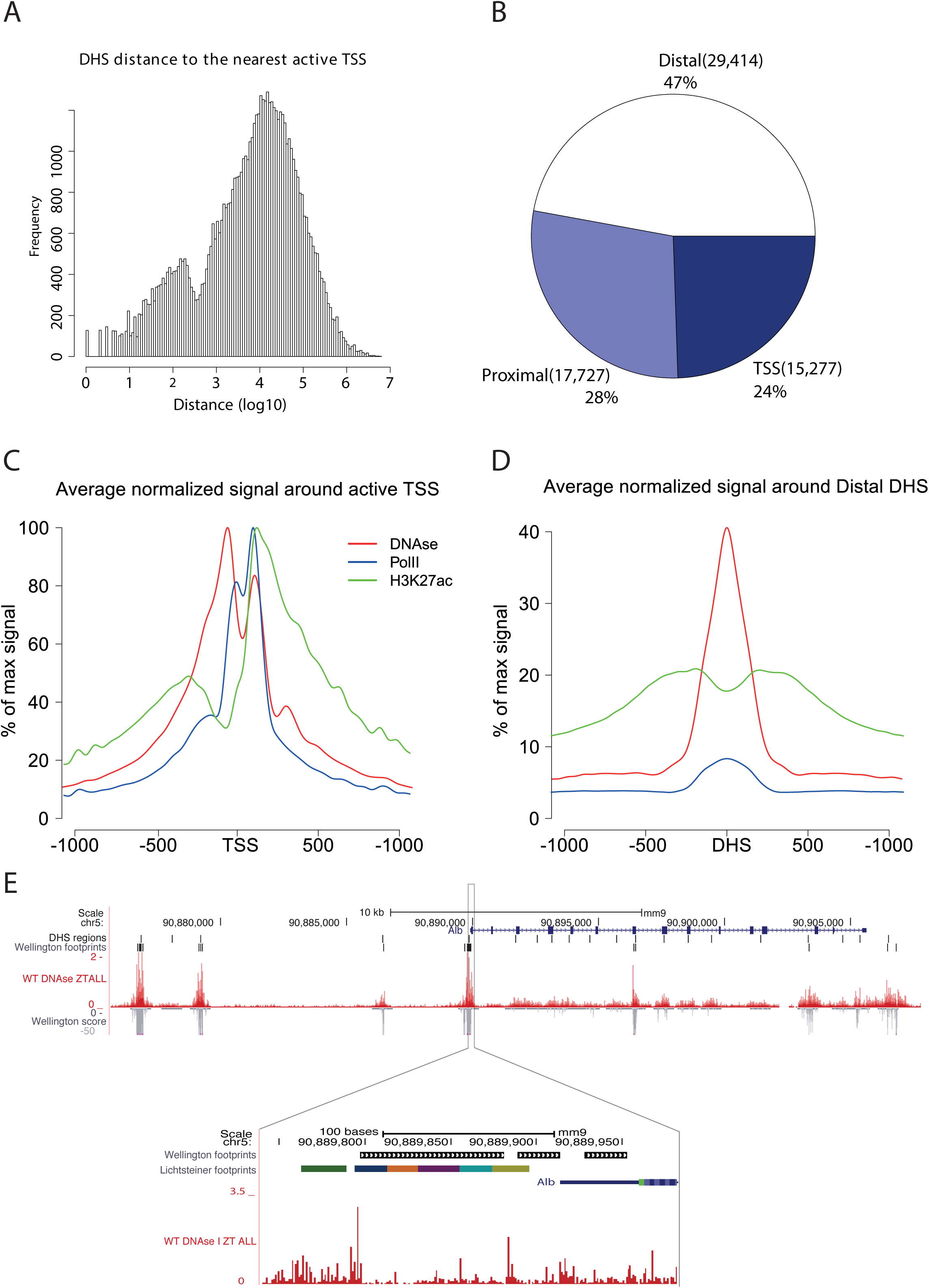
Characteristics of DHSs. A. Distribution of distances between DHSs and nearest active TSSs. We observe a bimodal distribution, with a first mode corresponding to DHSs in promoter regions (centered on 100 bp from the TSS) and a second mode centered on 10 kb from TSSs. B. Repartition of DHSs within three classes depending on their distance from the nearest TSS: 47% are more than 10 kb from a TSS and are classified as distal, 28% are between 1 kb and 10 kb away and are classified as proximal, and DHSs located 1 kb or less from a TSS represent 24% of all sites. C.-D. Pol II, DHS and H3K27ac signals around TSSs and distal DHSs (averages over all sites). Profiles were normalized so that the maximum around the TSS is 100%. E. DNase I signals (all time points are merged in the ZT All track) near the *Albumin* gene. Footprint detected using the Wellington algorithm are shown below the detected DHS sites. The promoter region is enlarged at the bottom, showing that the wide footprint detected in our data corresponds to previously established transcription factor binding sites (the colored boxed indicate protein complexes previously identified in [47]). Many sensitive regions locate din the gene body do not display footprints, probably due to high transcription of *Alb* in the liver.

**Fig S3.**
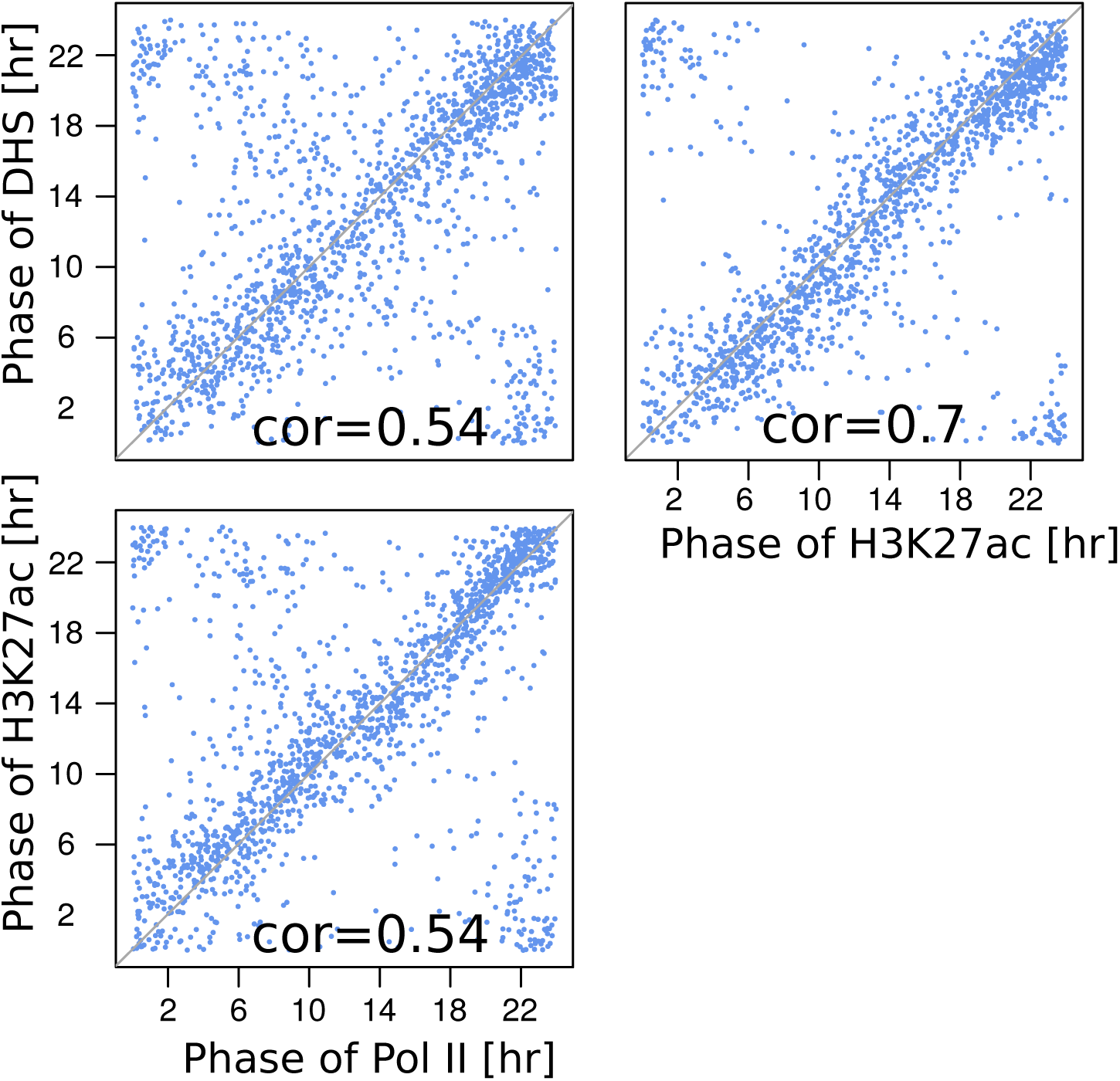
Phase relationships between DHS, Pol II, and H3K27ac at all DHS sites outside transcribed regions. Similarly to Figure 3D, high correlations and no phase shifts can still be observed outside of actively transcribed regions, demonstrating that this relationship is not only linked to active transcription.

**Fig S4.**
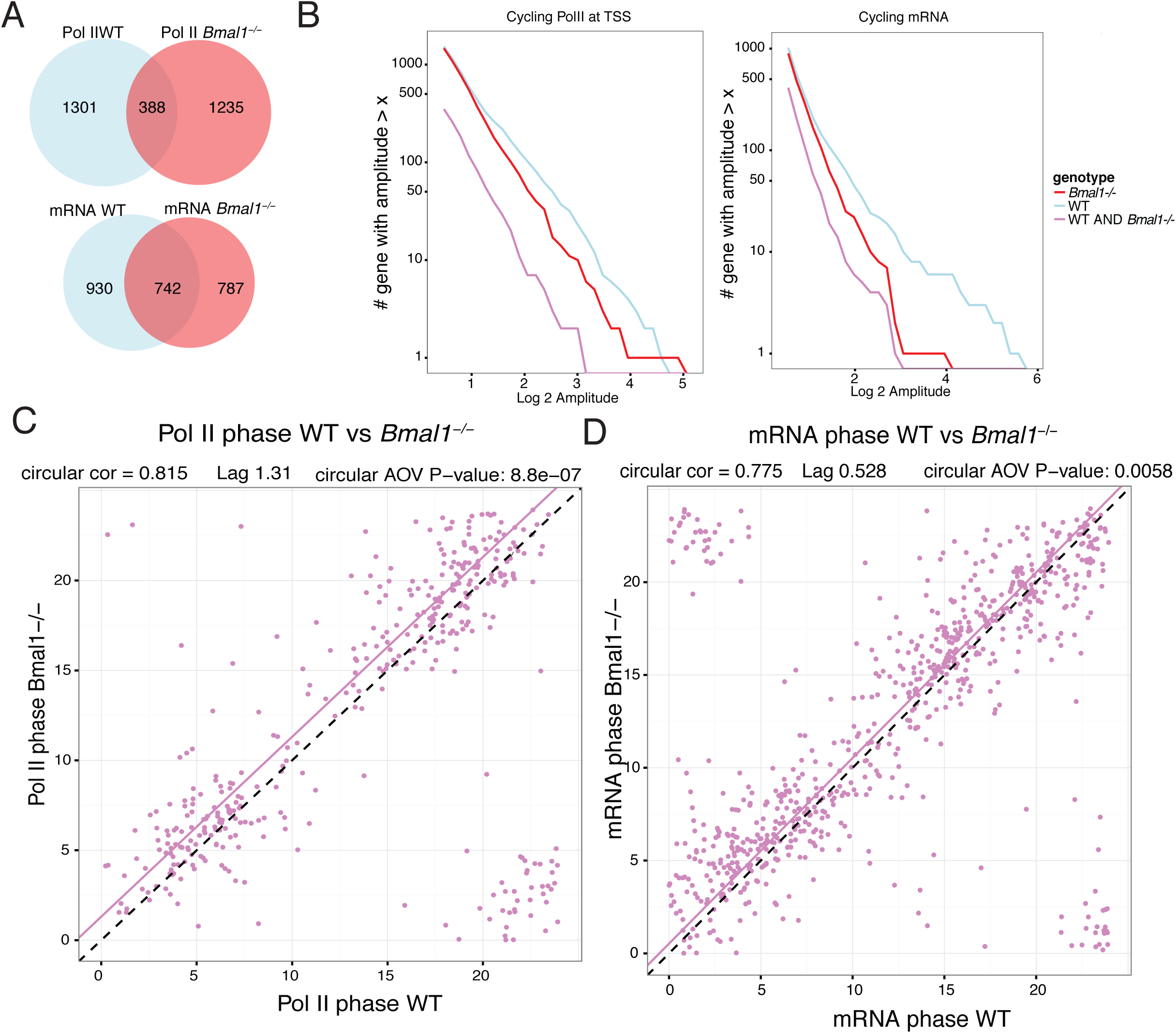
Diurnal oscillations in transcription and mRNA accumulation in WT and *Bmal1*^*-/-*^ livers. A. Number of oscillating genes in WT and in *Bmal1*^*-/-*^ mice using Pol II loadings at TSSs and mRNA. B. Cumulative count of oscillating genes (selected with p < 0.05, harmonic regression) in *Bmal1*^*-/-*^ and WT mice with log_2_ amplitude greater or equal than the values on the x-axis. Both Pol II loadings at TSSs and mRNA are shown. Values below 0.5 on the x-axis are not shown. C. Peak times (ZT times) of genes oscillating in WT and in *Bmal1*^*-/-*^ using Pol II loadings at TSS. D. Idem using mRNA accumulation profiles.

**Fig S5.**
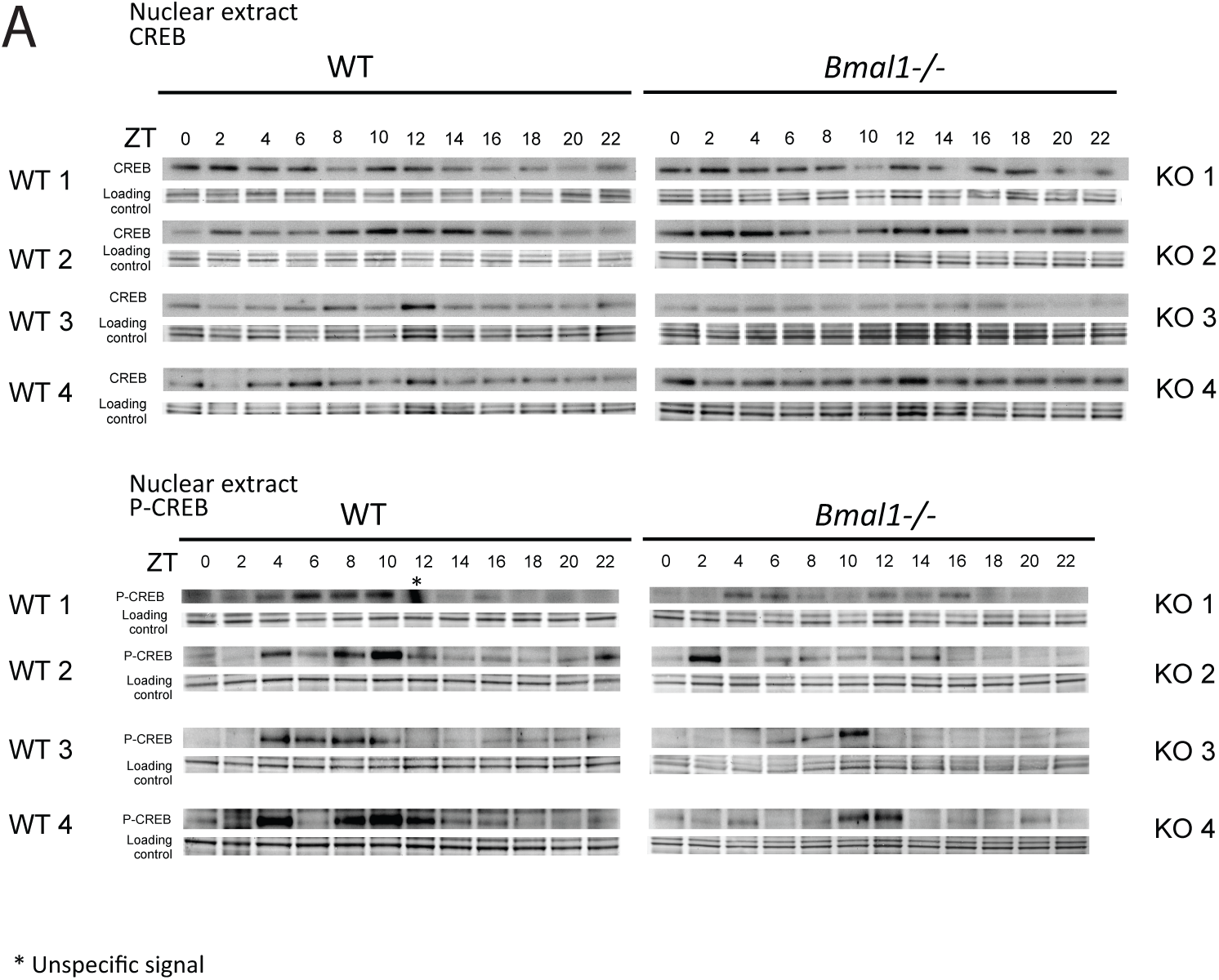
Western blot time-series of CREB and pCREB in nuclear extracts from WT and *Bmal1*^*-/-*^ livers (n=4 individual animals per time point). Quantifications and statistical analysis are shown in Figure 4. Loading control shows staining with naphtol blue black.

**Fig S6.**
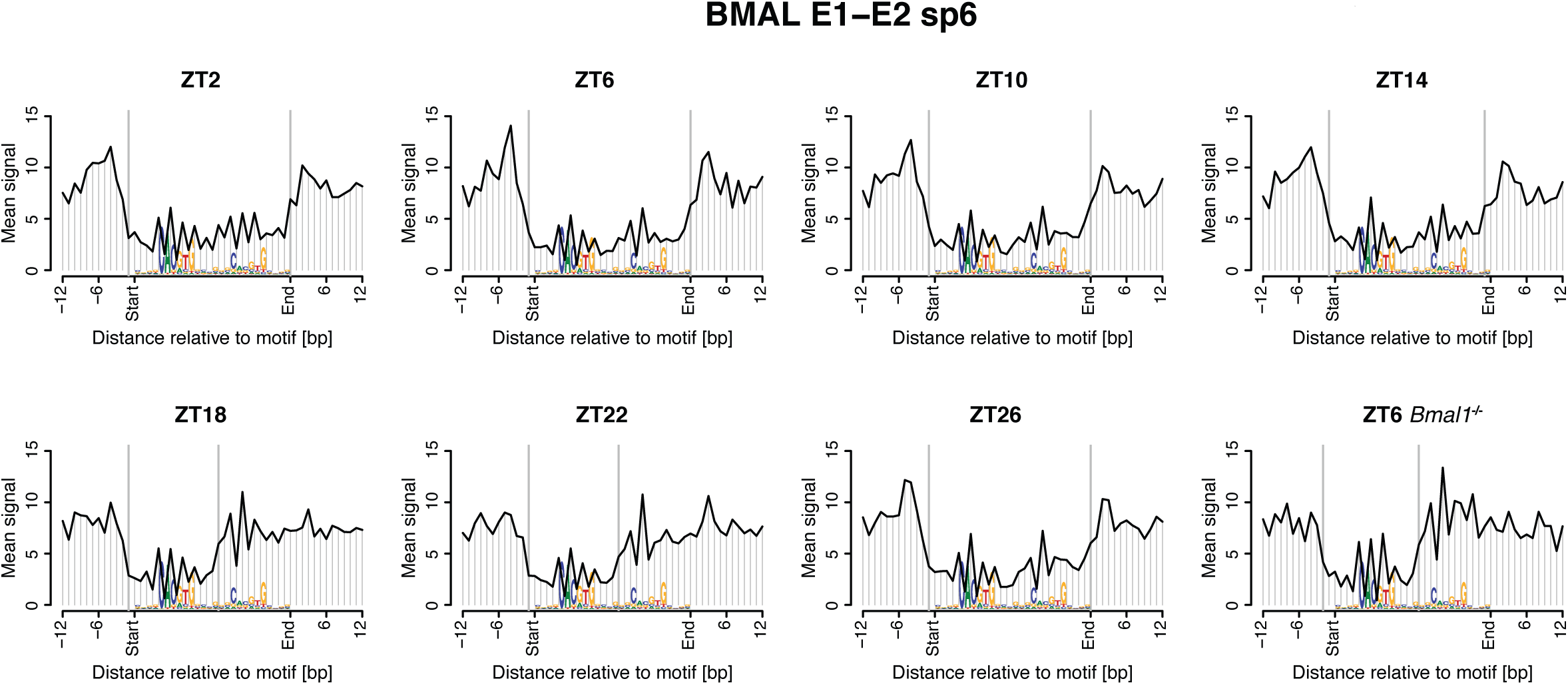
Genomic profiles of DNase I cuts around double E-boxes with a spacer of 6 bp (E1-E2 sp6) at all time points. The analysis is identical to that in Figure 6A. The analysis for ZT6 in *Bmal1*^*-/-*^ mice is also shown.

**Fig S7.**
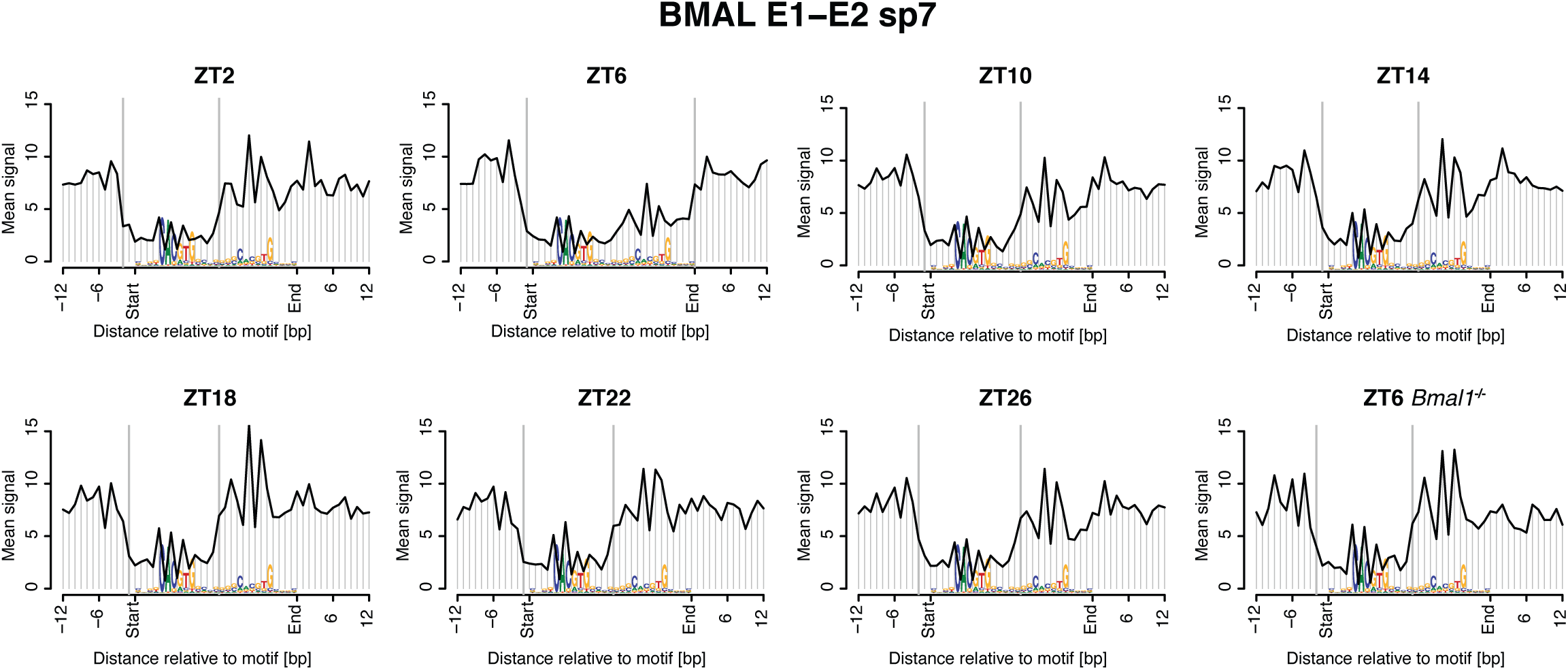
Idem as as Figure S6 but for double E-boxes with a spacer of 7 bp.

**Fig S8.**
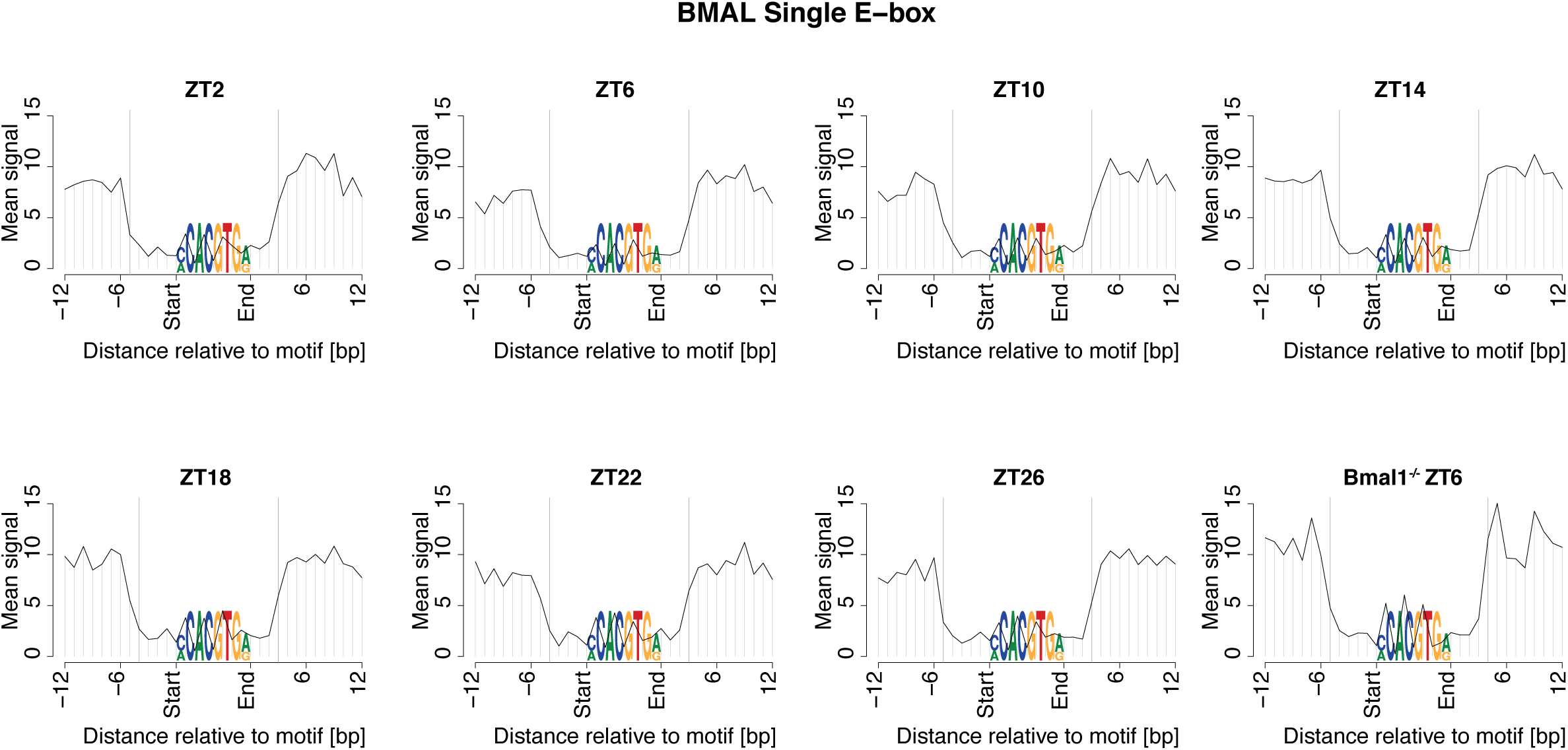
Idem as as Figure S6 but selecting BMAL1 bound DHSs containing single E-boxes. Otherwise the analysis is identical to Figures S6 and S7.

**Fig S9.**
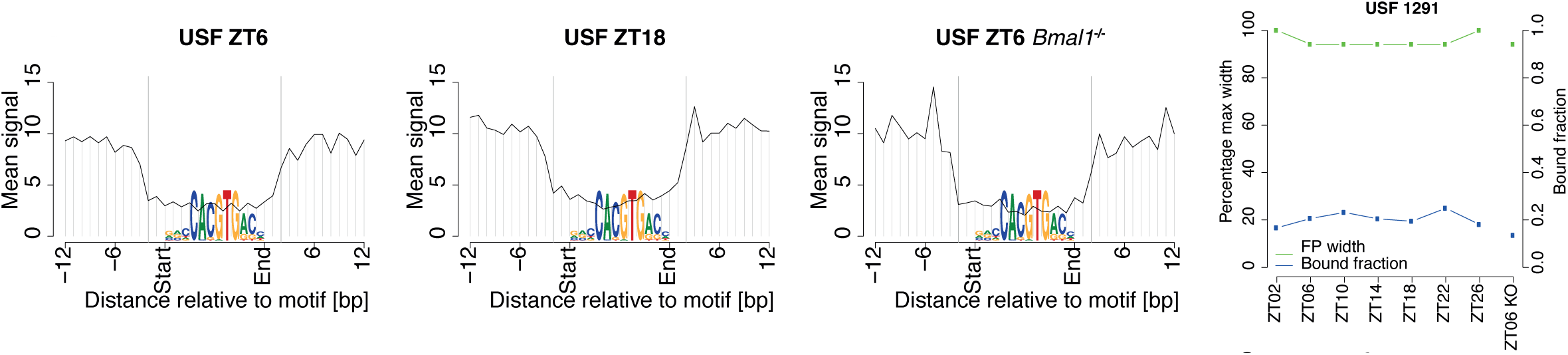
Idem as Figure 6A, but selecting DHSs bound by USF1 and containing a USF1 motif (E-box).

**Fig S10.**
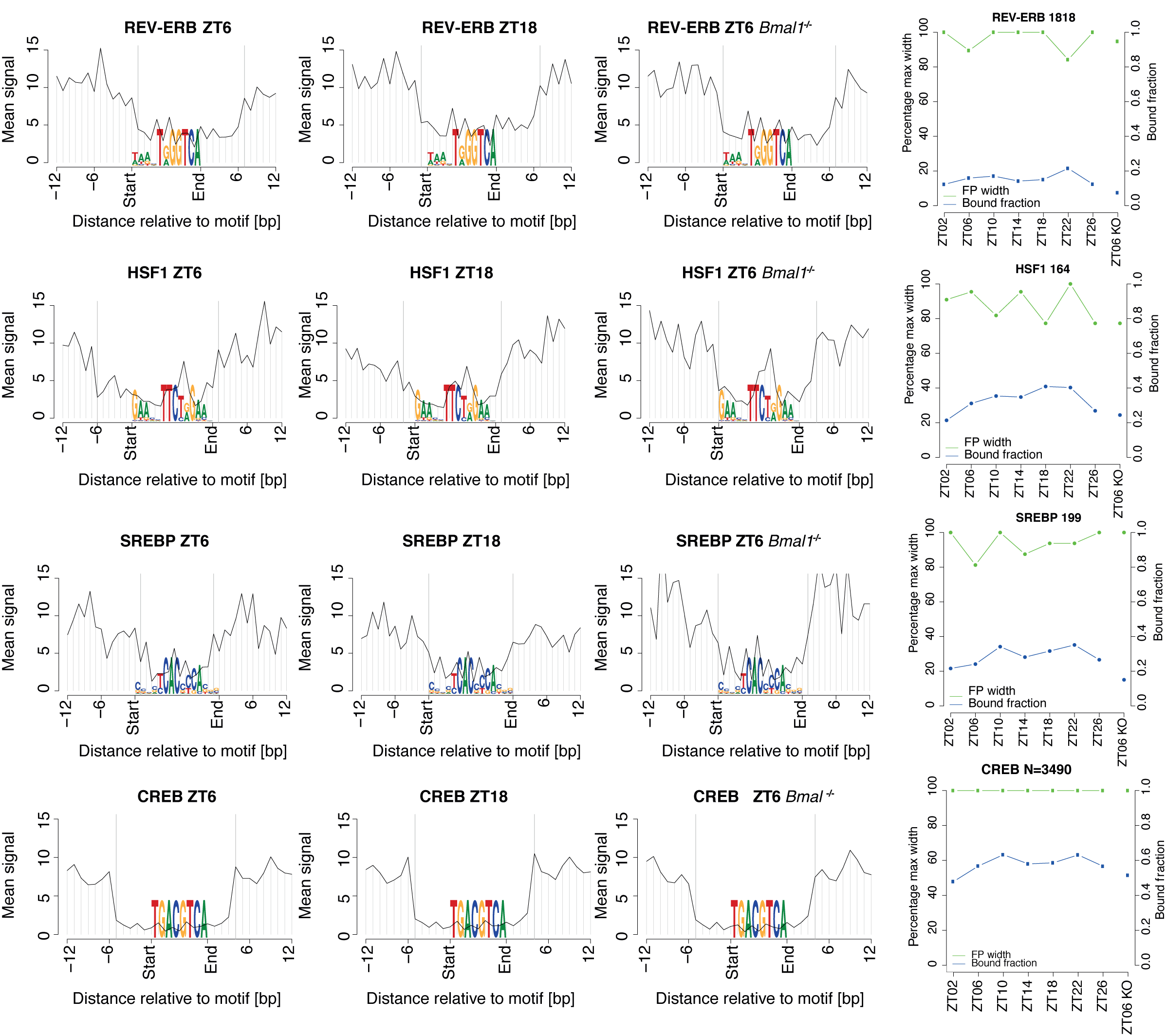
Idem as Figure 6A, but selecting DHSs bound by REV-ERB, HSF1, SREBP and CREB, and containing corresponding motifs. Here, DHS sites overlapped by a high ChIP-seq signal (Z score > 2) were considered.

**Supplementary Files**

**Table S1:** Quality control and mapping statistics for DNase I, Pol II and H3K27ac to the mm9 genome assembly.

**Table S2:** All identified DHSs with quantified signals for DNase I, Pol II and H3K27ac in WT and *Bmal1*^*-/-*^.

**Table S3:** Kegg and Reactome Pathway analysis of oscillating genes in mRNA accumulation in WT and *Bmal1*^*-/-*^ mice.

**Table S4:** Inferred activity (phase and amplitudes) for PWMs (DNA motifs) retained by the penalized generalized linear model using Pol II loadings at TSS and motif content in DHSs within 50kb from the gene TSSs. The consensus sequence, the source of the PWM, the number of targets and the sum of motifs in DHSs are listed.

**Movie S1:** Dynamics of DNase I, Pol II and H3K27ac at the *Dbp* locus. **Movie S2:** Dynamics of DNase I, Pol II and H3K27ac at the *Npas2* locus. **Movie S3:** 3D structure of the Hetero-tetramer of BMAL1/CLOCK (sp6). **Movie S4:** 3D structure of the Hetero-tetramer of BMAL1/CLOCK (sp7). **File S1**: Mixture model for DNase I-seq footprints.

**File S2:** Hetero-tetramer of BMAL1/CLOCK in .pdb format

## Acknowledgments

We thank Jacques Rougemont for insightful discussion on data analysis and interpretation, and Meini Busslinger for advice and protocols on DHS mapping. Computations and analyses were performed at the Vital-IT (http://www.vital-it.ch) center for high-performance computing of the Swiss Institute of Bioinformatics (SIB). High throughput sequencing was performed at the Lausanne Genomic Technologies Facility.

